# The brassinosteroid receptor gene *BRI1* safeguards cell-autonomous brassinosteroid signaling across tissues

**DOI:** 10.1101/2024.05.13.593848

**Authors:** Noel Blanco-Touriñán, Surbhi Rana, Trevor M. Nolan, Kunkun Li, Nemanja Vukašinović, Che-Wei Hsu, Eugenia Russinova, ChrisHan S. Hardtke

## Abstract

Brassinosteroid signaling is essential for plant growth as exemplified by the dwarf phenotype of loss-of-function mutants in *BRASSINOSTEROID INSENSITIVE 1 (BRI1)*, a ubiquitously expressed Arabidopsis brassinosteroid receptor gene. Complementation of brassinosteroid-blind receptor mutants by *BRI1* expression with various tissue-specific promoters implied that local brassinosteroid signaling may instruct growth non-cell-autonomously. Here we performed such rescues with a panel of receptor variants and promoters, in combination with tissue-specific transgene knockouts. Our experiments demonstrate that brassinosteroid receptor expression in several tissues is necessary but not sufficient for rescue. Moreover, complementation with tissue-specific promoters requires the genuine *BRI1* gene body sequence, which confers ubiquitous expression of trace receptor amounts that are sufficient to promote brassinosteroid-dependent root growth. Our data, therefore, argue for a largely cell-autonomous action of brassinosteroid receptors.

## INTRODUCTION

Brassinosteroids, such as the prototypical brassinolide, are endogenous key regulators of plant growth (Nomura et al., 2005, Roh et al., 2020). In Arabidopsis (*Arabidopsis thaliana*), brassinolide is sensed by the extracellular domains of the receptor kinases BRASSINOSTEROID-INSENSITIVE 1 (BRI1) and its homologs BRI1-LIKE 1 (BRL1) and BRI1-LIKE (BRL3) (Cano-Delgado et al., 2004, Hothorn et al., 2011, Li and Chory, 1997). Brassinolide binding promotes their interaction with co-receptors, triggering a phospho-transfer cascade that permits nuclear accumulation of downstream transcription factors to regulate target genes (Kim and Russinova, 2020, Wang et al., 2002, Yin et al., 2002, Zhu et al., 2013). These include brassinosteroid biosynthesis pathway genes, establishing a feedback loop that maintains brassinosteroid signaling homeostasis. Loss-of-function mutations in *BRI1* or genes encoding key brassinosteroid biosynthetic enzymes lead to severe growth retardation, including strongly impaired root growth (Choe et al., 1998, Gonzalez-Garcia et al., 2011, Hacham et al., 2011, Kang et al., 2017, Nomura et al., 2005, Oh et al., 2020, Szekeres et al., 1996). The root phenotype of brassinosteroid-related mutants presents various aspects (Oh et al., 2020). Although cells are generally shorter, this does not scale with the overall reduction in root growth because of the additional impact of brassinosteroid signaling on cell proliferation (Gonzalez-Garcia et al., 2011, Hacham et al., 2011, Clark et al., 2021). Furthermore, brassinosteroid signaling restricts formative divisions, mostly in the stele (Fridman et al., 2021, Graeff et al., 2021, Kang et al., 2017, Ohashi-Ito et al., 2023), and is also required for the proper formation of vascular tissues (Cano-Delgado et al., 2004, Holzwart et al., 2018, Kang et al., 2017, Tamaki et al., 2020, Yamamoto et al., 2001) which are in turn important for root growth maintenance (Fukuda and Ohashi-Ito, 2019, Hardtke, 2023). Recent morphometric 3D single-cell analyses found that brassinosteroid signaling promotes cellular anisotropy but is not required for volumetric cell growth (Fridman et al., 2021, Graeff et al., 2021) and also enforces accurate cell division plane orientation (Graeff et al., 2021). Moreover, single-cell RNA sequencing (scRNAseq) suggests that overall specification and development of the different root tissue layers progresses correctly in brassinosteroid receptor mutants (Graeff et al., 2021, Nolan et al., 2023). Thus, their reduced root growth can be parsimoniously explained by the combined effects of reduced cellular anisotropy and aberrant cell divisions (Fridman et al., 2021, Graeff et al., 2020, Kang et al., 2017). These effects may be aggravated by brassinosteroid-dependent inter-cell layer communication (Fridman et al., 2021, Vragovic et al., 2015), which is also a salient feature of wildtype root development since rate-limiting brassinosteroid biosynthesis genes are expressed in specific cell layers along a spatiotemporal gradient (Vukasinovic et al., 2021). Differential biosynthesis may thus be responsible for differential brassinosteroid effects along the root, with lower levels favoring meristematic activity and higher levels favoring cellular anisotropy (Vukasinovic et al., 2021).

The precisely aligned cell files that differentiate into the distinct root tissues are produced by apical stem cells at the root meristem tip and give rise to a stereotypic pattern of radial symmetry in the outer tissue layers and bilateral symmetry in the vascular cylinder (Fig. 1A). Intriguingly, although *BRI1* is expressed throughout the root (Fig. S1), the phenotype of *bri1* mutants can be complemented by transgenic expression of BRI1 under control of tissue-specific promoters (Hacham et al., 2011, Hategan et al., 2014, Kang et al., 2017, Vragovic et al., 2015). This could also be observed in *bri1 brl1 brl3 (bri*^*3*^) triple receptor null mutants (Kang et al., 2017, Vragovic et al., 2015) in which aspects of the *bri1* phenotype are aggravated (Cano-Delgado et al., 2004, Kang et al., 2017, Zhou et al., 2004), ruling out compensatory effects by *BRL1* or *BRL3* which are expressed at much lower levels than *BRI1* (Cano-Delgado et al., 2004, Nolan et al., 2023) (Fig. S1). To some degree, the extent of complementation depends on the tissue and the expression level. For example, excess epidermal BRI1 mimics brassinosteroid signaling gain-of-function effects (Fridman et al., 2014, Hacham et al., 2011, Vragovic et al., 2015), whereas high BRI1 levels in the vascular cylinder can trigger supernumerary formative divisions (Fridman et al., 2021, Graeff et al., 2021, Hategan et al., 2014, Kang et al., 2017). Also, more restricted *BRI1* expression in the two developing protophloem sieve element cell files can largely restore *bri*^*3*^ root growth but confers an intermediate rescue of cellular features (Graeff et al., 2020, Graeff et al., 2021, Kang et al., 2017). One key conclusion from these experiments is that brassinosteroid receptors may instruct growth non-cell-autonomously (Graeff et al., 2020, Hacham et al., 2011, Hategan et al., 2014, Savaldi-Goldstein et al., 2007, Vragovic et al., 2015, Fridman et al., 2014, Kang et al., 2017).

**Fig. 1.**
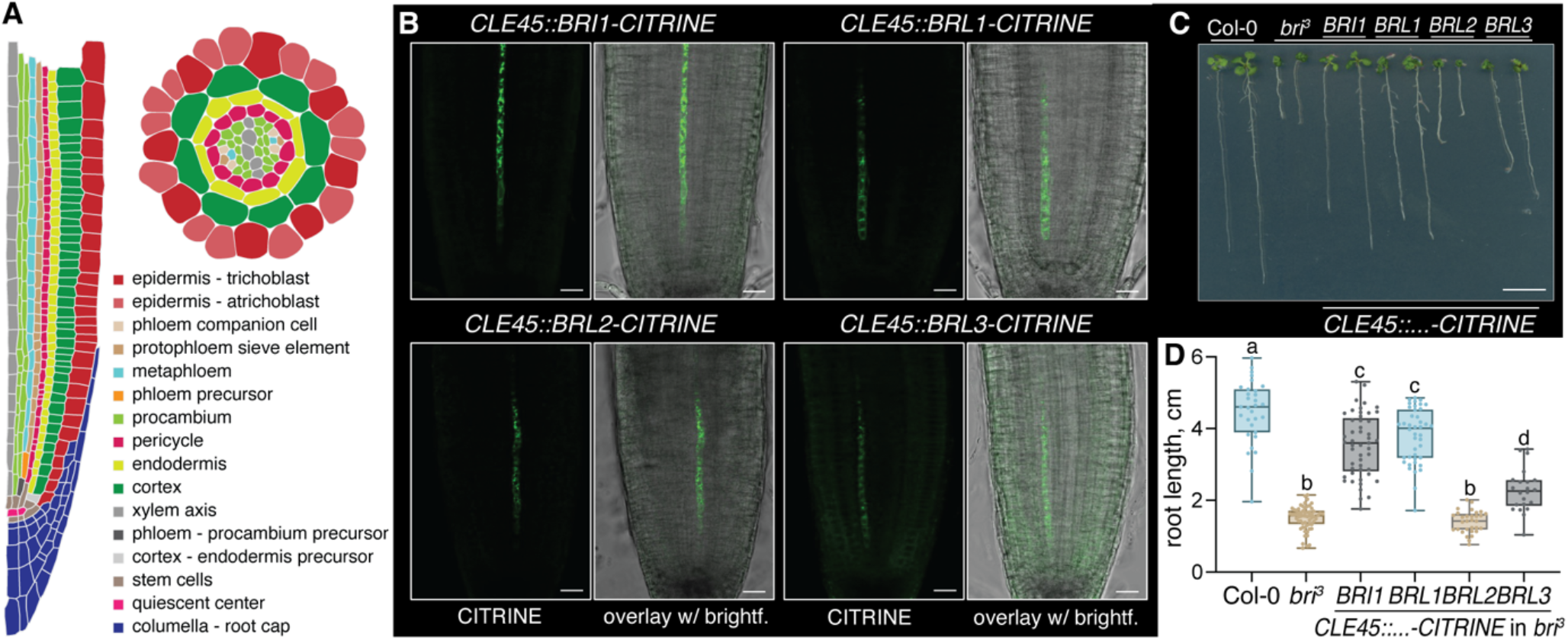
Arabidopsis brassinosteroid receptors are interchangeable in (partial) complementation of brassinosteroid-blind mutants with tissue-specific promoters. **(A)** Schematic presentation of an Arabidopsis root tip with a (half) longitudinal and a cross section. **(B)** Confocal microscopy imagesof root meristems from indicated genotypes (without counterstaining), showing CITRINE signal of indicated fusion proteins (green fluorescence, left panels)and their overlay with brightfield images (right panels). **(C)** Representative 9-day-old Col-0 wildtype, *bri*^*3*^ triple mutant and transgenic *bri*^*3*^ seedlingsexpressing indicated brassinosteroid receptor CITRINE fusion proteins under control of the CLE45 promoter. **(D)** Root growth quantification for 8-d-oldseedlings of the indicated genotypes. Box plots display 2nd and 3rd quartiles and the median, bars indicate maximum and minimum. Statistically significantdifferences (lower case letters) were determined by ordinary one-way ANOVA, p<0.004. Size bars are 20 μm in (B) and 1cm in (C).

Here, we sought to determine the nature of the proposed non-cell-autonomous *BRI1* effects. We found that the *BRI1* gene promotes its own ubiquitous expression through gene body-intrinsic sequences and that trace amounts of BRI1 across multiple tissues are necessary to complement *bri*^*3*^ mutants. Thus, our data suggest that the brassinosteroid receptor acts largely cell-autonomously.

## RESULTS

### Brassinosteroid receptor genes are interchangeable in bri^3^ complementation

We first evaluated whether *BRI1* homologs can reproduce the reported *bri*^*3*^ rescue observed upon expression of BRI1-CITRINE fusion protein with the phloem-specific *COTYLEDON VASCULAR PATTERN 2 (CVP2), BARELY ANY MERISTEM 3 (BAM3)* and *MEMBRANE-ASSOCIATED KINASE REGULATOR 5 (MAKR5)* promoters (Graeff et al., 2020, Kang et al., 2017). Because we previously found that the extent of rescue by *CVP2::BRI1-CITRINE* depends on transgene expression level and dosage (Graeff et al., 2020), we expressed the receptor fusion proteins using the promoter of the protophloem sieve element-specific gene, *CLAVATA3/EMBRYO SURROUNDING REGION 45 (CLE45)* (Kang and Hardtke, 2016, Rodriguez-Villalon et al., 2014), which is expressed at higher levels than *CVP2* (Fig. S1). Substantial *bri*^*3*^ rescue was observed with BRI1, BRL1 and BRL3, but not with the related (non-brassinosteroid) receptor BRL2 (Fig. 1B-D). Consistent with previous results (Cano-Delgado et al., 2004, Zhou et al., 2004), our findings reiterate that BRI1, BRL1 and to some extent BRL3 are functionally equivalent in their ability to rescue the root growth of *bri*^*3*^ when expressed at similar levels.

### Brassinosteroid signaling in multiple tissues is necessary for comprehensive bri^3^ rescue

The epidermis is a key tissue for brassinosteroid perception (Hacham et al., 2011, Savaldi-Goldstein et al., 2007), and consistently tissue-specific CRISPR/Cas9-mediated *BRI1* transgene knockout in the epidermis using the *WEREWOLF (WER)* promoter reverts the phenotype of complemented *bri1* single mutants (Nolan et al., 2023). This contrasts with *bri*^*3*^ rescue by expression of brassinosteroid receptors with phloem-specific promoters. To directly test the impact of phloem-specific BRI1 dosage on *bri*^*3*^ rescue, we expressed Cas9 under the control of the *SHORT ROOT (SHR)* promoter together with *BRI1*-specific single guide RNAs (Nolan et al., 2023) *(SHR::Cas9*^*BRI1*^), in a *CVP2::BRI1-CITRINE* reference line that contained three concatenated transgenes (Graeff et al., 2020). *SHR* is expressed in the vascular cylinder except in the developing protophloem (Fig. S1) (Kang et al., 2017, Kim et al., 2020). However, because the *SHR* promoter is active in all vascular stem cells including phloem precursors (Kang et al., 2017, Kim et al., 2020), we could recover transformants in which BRI1-CITRINE signal was no longer detectable in the phloem (Fig. S2A). These plants did not display growth defects to the same extent as *bri*^*3*^ (Fig. S2B). To corroborate this finding, we transformed the *SHR::Cas9*^*BRI1*^ construct into another homozygous *bri*^*3*^ rescue line that expressed BRI1-CITRINE under the control of the phloem pole-specific *MAKR5* promoter (Fig. S1) (Kang and Hardtke, 2016). In the progeny of several independent lines that segregated the *SHR::Cas9*^*BRI1*^ transgene, we could compare siblings in which the phloem pole BRI1-CITRINE signal was present with those in which it was absent. Again, the latter did not display a *bona fide bri*^*3*^ phenotype (Fig. 2A), however compared to their siblings, root growth was reduced to an intermediate length by on average c. 31% (Fig. 2B, Fig. S2C). Interestingly, the *MAKR5::BRI1-CITRINE* rescue lines also displayed faint yet readily detectable plasma-membrane-localized fluorescent signal in the epidermal tissues that was distinct from background fluorescence (Fig. 2C) and persisted in the presence of the *SHR::Cas9*^*BRI1*^ transgene (Fig. 2D; Fig. S2D-E). To determine whether this signal originated from the *MAKR5::BRI1-CITRINE* transgene and whether it had an impact on root growth, we also transformed a *WER::Cas9*^*BRI1*^ construct. *WER* is expressed in epidermal tissues (Fig. S1) (Lee and Schiefelbein, 1999, Nolan et al., 2023). In the progeny of the *WER::Cas9*^*BRI1*^ plants, we frequently observed loss of the epidermal signal, indicating that it indeed originated from the *MAKR5::BRI1-CITRINE* transgene (Fig. 2E; Fig. S2F). Moreover, in such seedlings root growth complementation was lost and the plants resembled *bri*^*3*^ mutants (Fig. 2A-B; Fig. S2C) despite the continued presence of BRI1-CITRINE signal in the phloem poles (Fig. 2E). Collectively these results indicate that although the phloem contributes to brassinosteroid-mediated root growth, additional low levels of BRI1-CITRINE expression in the epidermis are accountable for the comprehensive rescue of *bri*^*3*^ mutants.

**Fig. 2.**
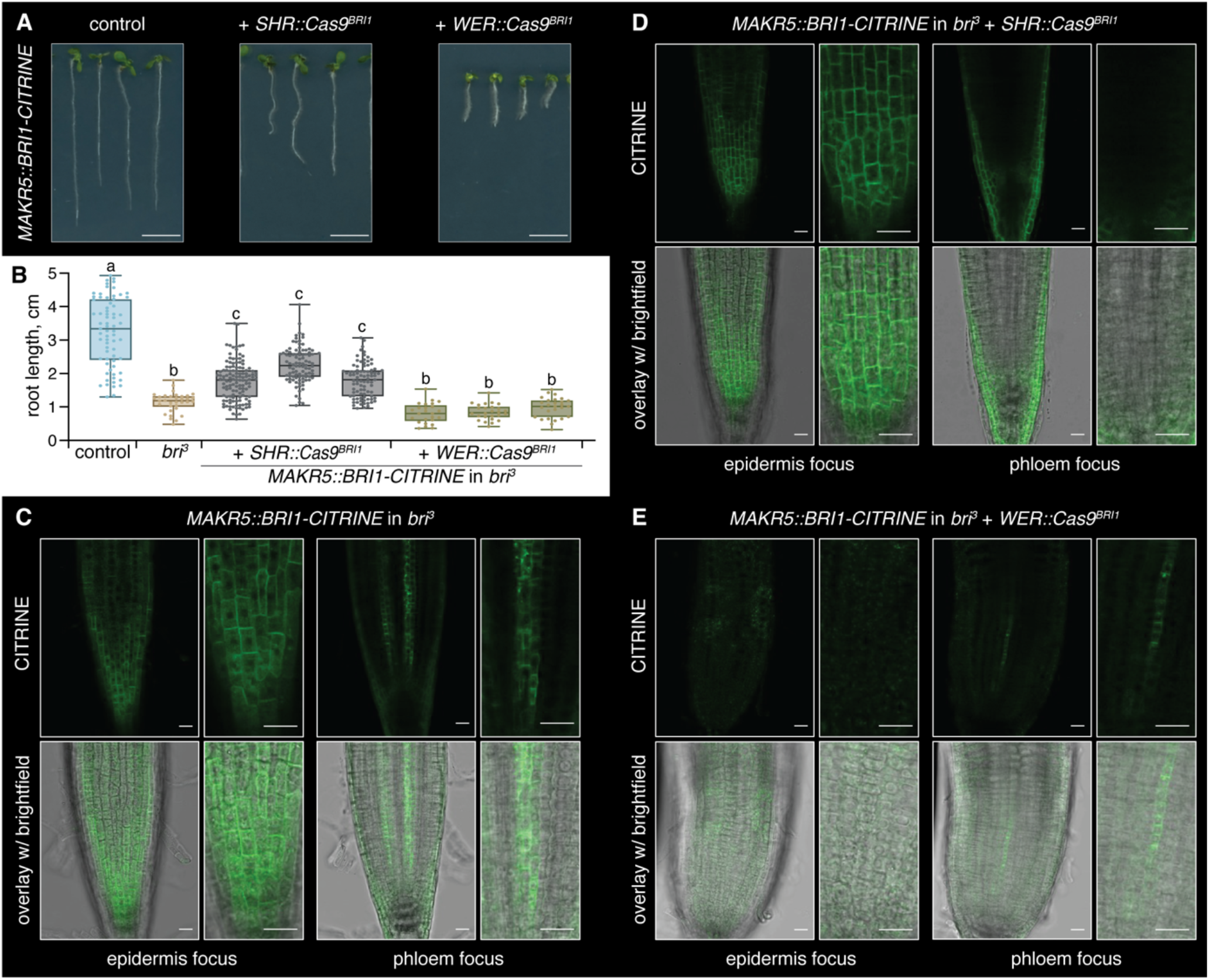
Brassinosteroid perception in both phloem and epidermis is necessary for comprehensive rescue of *bri*^3^ mutants. **(A)** Representative 8-day-old *bri*^3^ seedlings complemented with a *MAKR5::BRI1 –CITRINE* transgene (left) and the same line combined with tissue-specific CRISPR/Cas9 *BRI1*knockout using the *SHR* (middle) or *WER* (right) promoters. **(B)** Root growth quantification for 8-d-old seedlings of the indicated genotypes, threeindependent CRISPR/Cas9 knockout lines are shown. Box plots display 2nd and 3rd quartiles and the median, bars indicate maximum and minimum.Statistically significant differences (lower case letters) were determined by ordinary one-way ANOVA, p<0.001. **(C-E)** Confocal microscopy images of rootmeristems from indicated genotypes imaged with relaxed settings and without any counterstaining. BRI1-CITRINE signal (green fluorescence) is observedin the epidermis although the *MAKR5* promoter is phloem pole-specific (C). CRISPR/Cas9 knockout using either the stele-specific *SHR* or the epidermis-specific *WER* promoter leads to disappearance of phloem pole (D) or epidermal signal (E), respectively. Size bars are 20 μm in microscopy images and1cm in seedling images.

### BRI1 transgenes display weak epidermal signal independent of tissue-specific promoters

Considering the importance of epidermal *BRI1* expression in the *MAKR5::BRI1-CITRINE* line, we next monitored various other *bri*^*3*^ rescue lines. In the *CVP2::BRI1-CITRINE* lines, epidermal signal was difficult to detect even if BRI1-CITRINE fluorescence in the protophloem was strong. However, when imaged without counterstaining and depending on the confocal microscopy instrument, faint plasma-membrane-localized signal that was absent from background controls could be detected (Fig. 3A-B). Such signal was more readily detected in *CLE45::BRI1-CITRINE* and *BAM3::BRI1-CITRINE* seedlings (Fig. 3C) as well as *SHR::BRI1-GFP* seedlings (Fig. 3D), indicating that it did not depend on the fluorophore. Consistently, it was also observed in newly generated *SHR::BRI1-CITRINE* seedlings (Fig. S2G). Moreover, the epidermal signal did not depend on the genetic background (Fig. 3E, Fig. S3A). Introduction of the *WER::Cas9*^*BRI1*^ construct into *CLE45::BRI1-CITRINE* seedlings again triggered reversion to a *bri*^*3*^ phenotype despite continued signal in the phloem (Fig. 3F). These experiments corroborated that the faint epidermal expression was essential for mutant complementation, although it was considerably weaker than the epidermal BRI1-GFP signal observed upon expression with the atrichoblast-specific *GLABRA 2 (GL2)* promoter or the native *BRI1* promoter (Fig. 3G, Fig. S1). To further pinpoint the origin of the epidermal signal, we monitored nuclear localized NLS-SCARLET fusion protein expressed under control of the *CLE45* promoter. NLS-SCARLET was exclusively detected in developing protophloem sieve elements, both when imaged alone or in *CLE45::BRI1-CITRINE* background (Fig. 3H, Fig. S3B). These findings suggest that the observed epidermal BRI1-CITRINE signal was not due to the promoters, the fluorophores or regulatory elements in the T-DNA vectors.

**Fig. 3.**
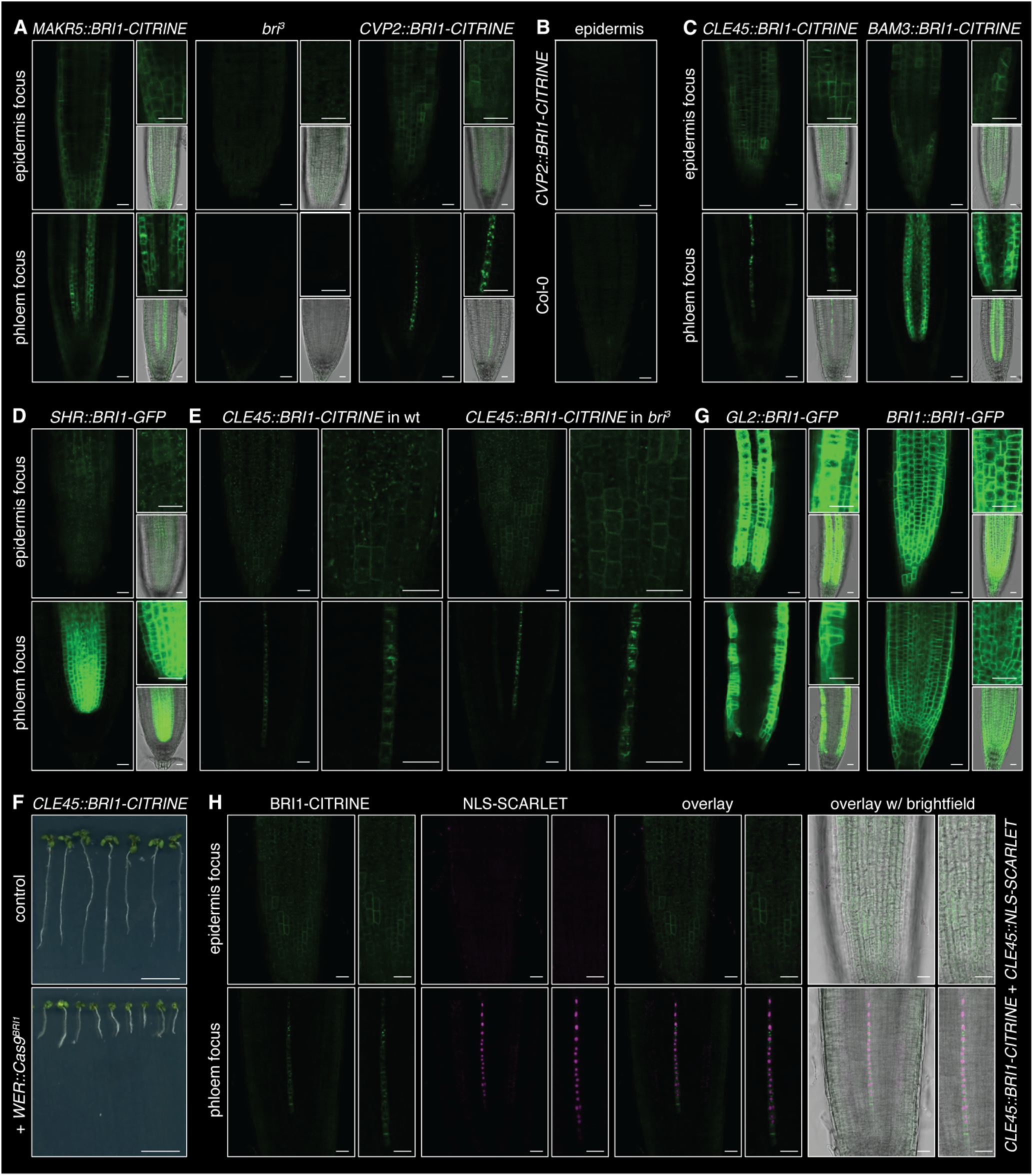
The *BRI1* gene body sequence confers low level ubiquitous expression of trace receptor protein amounts. **(A-D)** Confocal microscopy images of root meristems from seedlings expressing BRI1-CITRINE or –GFP fusion protein under control of different tissue-specific promoters in *bri*^*3*^background. Note the faint plasma-membrane-localized signal that is absent in *bri*^*3*^ controls. **(E)** Comparison of phloem sieve element-specific *CLE45*promoter-driven BRI1-CITRINE signal in morphologically wild type (*bri1* +/− *brl1* +/− *brl3*+/−) and *bri*^*3*^ background. **(F)** Representative 8-day-old *bri*^*3*^ seedlings complemented with a *CLE45::BRI1 –CITRINE* transgene (top) and the same line combined with tissue-specific CRISPR/Cas9 *BRI1* knockout using the *WER* promoter (bottom). **(G)** Confocal microscopy images of root meristems from seedlings expressing BRI1-GFP fusion protein under control of the atrichoblast-specific *GL2* or the native *BRI1* promoter in *bri*^*3*^ background. **(H)** Confocal microscopy images of a root meristem expressing both BRI1-CITRINE fusion protein (green fluorescence) and nuclear localized NLS-SCARLET protein (magenta fluorescence) under control of the *CLE45* promoter. Note exclusively phloem sieve element-specific NLS-SCARLET signal. Size bars are 20 μm in microscopy images and 1cm in seedling images.

### Ectopic BRI1 transgene expression is not a result of brassinosteroid feedback regulation

Matching their capacity to rescue *bri*^*3*^, faint epidermal signal was also detected in *CLE45::BRL1-CITRINE* and *CLE45::BRL3-CITRINE* plants but not in *CLE45::BRL2-CITRINE* plants (Fig. S4A-C). This raised the question whether the ectopic signal may result from auto-regulatory feedback, since *bri*^*3*^ rescue by BRI1 depends on brassinosteroid biosynthesis (Graeff et al., 2020). To test this idea, we first treated *BRI1-CITRINE* lines with the brassinosteroid biosynthesis inhibitor, brassinazole. This abolished *bri*^*3*^ rescue although the ectopic BRI1-CITRINE expression persisted (Fig. S4D-E). Oppositely, brassinolide treatment did not affect the epidermal signal (Fig. S4F). We also found a hypomorphic mutation (D48del) in the brassinosteroid biosynthetic enzyme DWARF1 (DWF1) (Choe et al., 1999) that reverts the *bri*^*3*^ rescue in *CVP2::BRI1-CITRINE* background but can be restored by brassinolide application (Fig. S4G). Epidermal BRI1-CITRINE signal in this line also did not respond to external supply of brassinolide (Fig. S4H). Moreover, *BRI1* control constructs with a point mutation that abolishes BRI1 kinase activity (E1078K; *CLE45::BRI1*^*KDD*^*-CITRINE)* (Zhang et al., 2018) could not complement *bri*^*3*^ and still displayed epidermal signal (Fig. S5A-B). Together, these experiments verify that the ectopic transgene expression was not a result of restored brassinosteroid perception itself.

### Gene body-intrinsic DNA sequences drive trace ubiquitous BRI1 gene expression

In parallel, we tested whether BRI1 kinase activity is not only necessary but also sufficient to confer *bri*^*3*^ rescue. To this end we introduced a constitutively active chimeric receptor composed of the BAK1-INTERACTING RECEPTOR-LIKE KINASE 3 (BIR3) extracellular domain and the intracellular BRI1 kinase domain (BIR3^EXT^BRI1^INT^) (Hohmann et al., 2020) into the *bri*^*3*^ background. The BIR3^EXT^BRI1^INT^ chimera can trigger strong gain-of-function phenotypes that mimic brassinosteroid pathway hyperactivity (Hohmann et al., 2020). Expression of a BIR3^EXT^BRI1^INT^-CITRINE fusion under control of the *CLE45* promoter produced a range of phenotypes that were apparently related to transgene dosage and expression. While we noticed the described gain-of-function phenotype of short, twisted roots (Fig. S5A), largely rescued yet still twisting roots were also observed (Fig. S5C). However, in all seedlings, BIR3^EXT^BRI1^INT^-CITRINE fluorescence was observed throughout the root tissues (Fig. S5D-E). These experiments thus not only suggested that constitutively activated BRI1 kinase domain can rescue *bri*^*3*^, but also that it confers ectopic expression in tissues other than the epidermis. To verify this notion, we reevaluated the *CVP2::BRI1-CITRINE* reference line by scRNAseq compared to wildtype. Interrogation of these data with a dedicated *BRI1-CITRINE* reference sequence revealed low yet widespread expression of the *BRI1-CITRINE* transcript, contrasting with the more restricted expression of *CVP2* (Fig. 4A-B, tab. S1). Moreover, reanalysis of scRNAseq data from a *bri*^*3*^ line expressing a BRI1-GFP fusion protein under control of the *GL2* promoter (Nolan et al., 2023) yielded a similar result with a *BRI1-GFP* reference. I.e., besides in atrichoblasts as expected, *BRI1-GFP* expression was also observed in all other tissues (Fig. 4C-D, tab. S1).

**Fig. 4.**
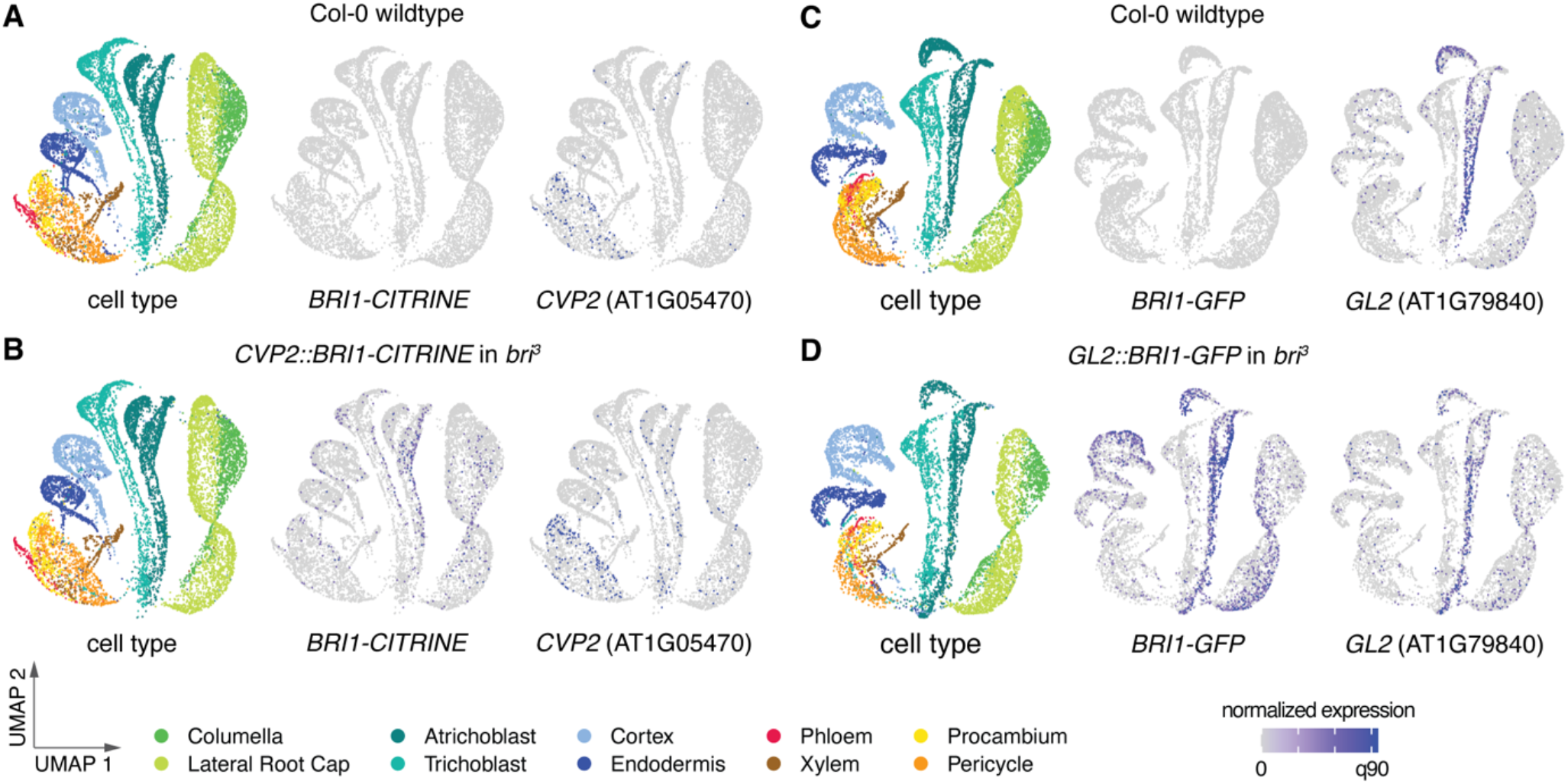
Below detection threshold levels of brassinosteroid receptor are ubiquitously expressed in brassinosteroid-blind mutants complemented with tissue-specific promoters. **(A-D**) UMAP representation of single cell RNA sequencing data obtained from Col-0 wild type or *bri*^*3*^ mutant root tips(partially) complemented with BRI1-CITRINE or BRI1-GFP fusion protein expressed with either the *CVP2* (A-B) or the *GL2* promoters (C-D). Left panels:cell type overviews, dots represent cells. Middle and right panels: Blue dots represent cells with target gene expression. Note the difference between detected transgene transcripts and endogenous wild type references.

Because brassinosteroid receptor genes in Arabidopsis as well as other species typically do not contain any introns (https://phytozome-next.jgi.doe.gov) (Navarro et al., 2015) we concluded that ectopic *BRI1* transgene expression may reflect an inherent feature of the *BRI1* coding sequence. To directly test this idea, we engineered a recoded version of *BRI1 (BRI1*^*REC*^) in which 907 of the 3,591 nucleotides were exchanged (Fig. S6). When BRI1^REC^-CITRINE fusion protein was expressed under the control of the *BRI1* promoter it complemented the *bri*^*3*^ mutant (Fig. 5A-D) similar to *BRI1:BRI1-CITRINE, BRI1:BRL1-CITRINE* or *BRI1:BRL3-CITRINE* controls (Fig. S7A-B), confirming BRI1^REC^ functionality. However, when expressed with phloem-specific promoters no rescue was observed (Fig. 5E, Fig. S7C), although BRI1^REC^-CITRINE was readily detectable in the phloem, whereas epidermal expression was absent (Fig. 5F-G, Fig. S7D-E). The observation that *bri*^*3*^ rescue by *CVP2::BRI1-CITRINE* is accompanied by resistance to external application of CLE45 peptide in a dosage-dependent manner (Graeff et al., 2020) gave us a handle to independently verify BRI1^REC^-CITRINE functionality in the phloem. CLE45-resistance was observed whenever *BRI1* was expressed under protophloem(pole)-specific promoters (Fig. S7F). Moreover, *MAKR5::BRI1-CITRINE bri*^*3*^ seedlings that carried the *SHR::Cas9*^*BRI1*^ construct and had lost BRI1-CITRINE signal in the phloem pole also had lost their CLE45-resistance (Fig. S7G), corroborating that it is a consequence of high BRI1 activity in the phloem (Breda et al., 2019, Graeff et al., 2020). Consistently, *bri*^*3*^ or wildtype seedlings that expressed a *MAKR5::BRI1*^*REC*^*-CITRINE* transgene displayed CLE45-resistance (Fig. S7H). In summary our data suggest that the *BRI1* gene body contains regulatory sequences that confer low level ubiquitous gene expression throughout the root and are lost in a recoded *BRI1* gene.

**Fig. 5.**
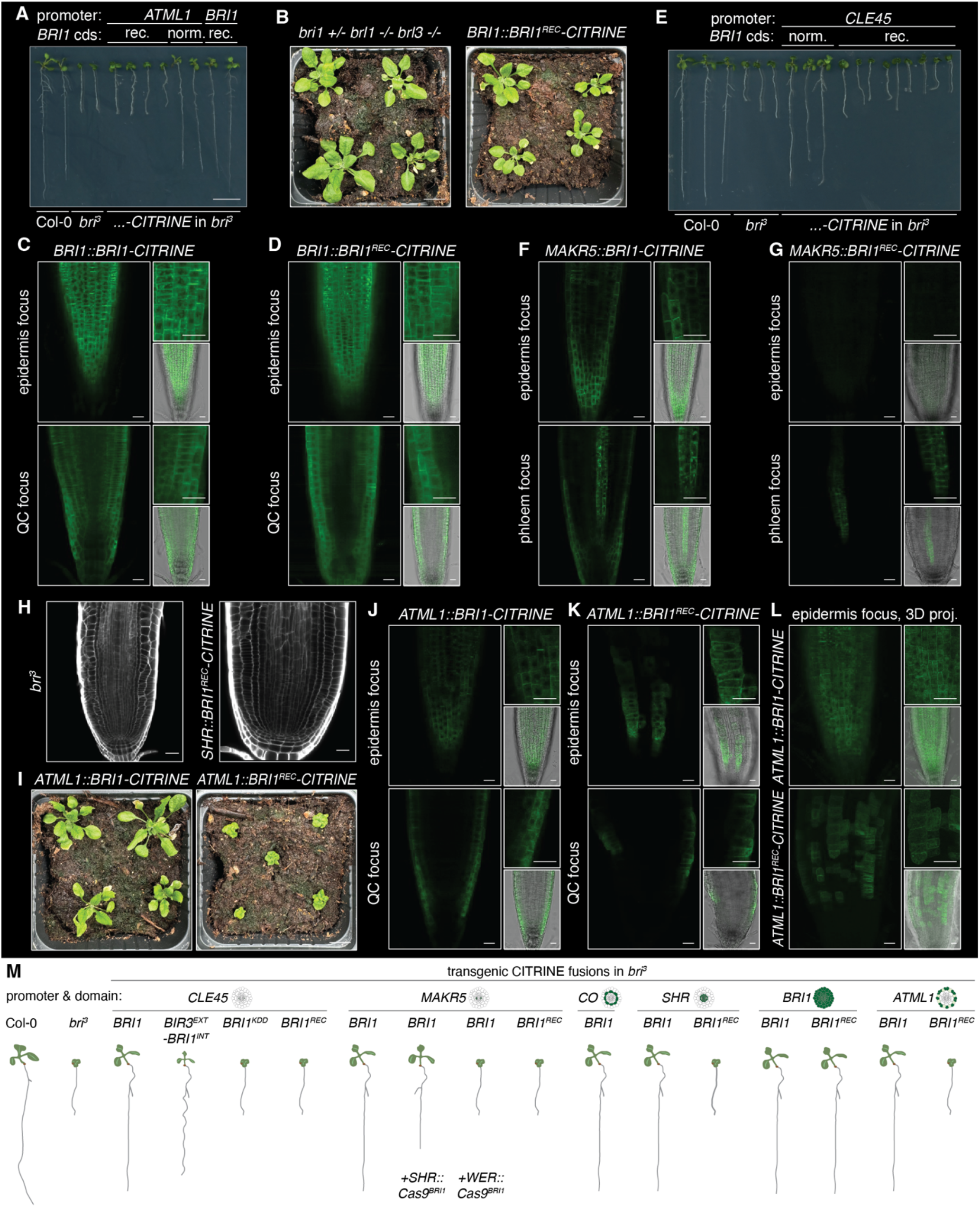
Ubiquitously expressed trace amounts of brassinosteroid receptor are necessary and sufficient to complement brassinosteroid-blind mutants. **(A)** Representative8-day-old seedlings carrying *BRI1-CITRINE* transgenes made with the genuine (norm.) *BRI1* coding sequence (cds) or a recoded version (rec.; c. 25% of nucleotides exchanged;*BRI1::BRI1*^*REC*^*-CITRINE*), compared to wildtype and *bri*^*3*^ controls. Note the difference in seedlings expressing either version under the epidermis-specific *ATML1* promoter. **(B)** 3-week-old *bri*^*3*^mutants complemented by a *BRI1::BRI1*^*REC*^*-CITRINE* transgene (right) compared to morphologically wild type *brl1 brl3* double mutants segregating from the transformed parental line (left). **(C-D)** Confocal microscopy images of root meristems from seedlings expressing BRI1 –CITRINE fusion proteins from the original or recoded *BRI1*cds under control of the native BRI1 promoter. **(E)** Representative 8-day-old seedlings expressing BRI1-CITRINE or BRI1^REC^-CITRINE fusion protein under control of the CLE45promoter in *bri*^*3*^ mutants, compared to wildtype and *bri*^*3*^ controls. **(F-G)** Confocal microscopy images of root meristems from seedlings expressing BRI1-CITRINE or BRI1^REC^-CITRINE fusion protein under control of the MAKR5 promoter in *bri*^*3*^ mutants, illustrating the absence of epidermal BRI1^REC^-CITRINE signal. **(H)** Confocal microscopy images ofcalcofluor-stained root meristems from *bri*^*3*^ mutants (left) and their counterparts expressing BRI1^REC^-CITRINE fusion protein under control of the *SHR* promoter (right). **(I)** 3-week-old brP mutants carrying an *ATML1::BRI1-CITRINE* (left) or *ATML1::BRI1*^*REC*^*-CITRINE* transgene (right). Note the difference compared to the complementation obtained with a *BRI1::BRI1*^*REC*^*-CITRINE* transgene (B). **(J-L)** Confocal microscopy images of root meristems from the indicated genotypes. QC = quiescent center. 3D projections focused on theepidermis are shown in (L). **(M)** Schematic overview of BRI1-related transgenic lines produced in this study and their representative phenotypes. Size bars are 20 μm in microscopyimages and 1cm in seedling images.

### Brassinosteroid signaling in individual tissues is not sufficient for bri^3^ complementation

Whether trace amounts of BRI1 in tissues other than the epidermis and phloem vasculature are also required for comprehensive *bri*^*3*^ rescue was difficult to determine but appeared likely. For instance, CRISPR/Cas9-mediated *BRI1* knockout in the cortex cell layer was recently shown to largely revert complementation of *bri1* single mutants (Nolan et al., 2023). Oppositely, expression of a BRI1-GFP fusion protein with a cortex-specific *(COR*; AT1G09750) promoter complemented *bri*^*3*^ root growth (Fig. S8A), but consistently these seedlings also showed epidermal BRI1-GFP signal (Fig. S8B). To determine whether brassinosteroid signaling in any individual tissue is sufficient to rescue the root growth of *bri*^*3*^, we expressed the BRI1^REC^ variant with additional promoters. BRI1^REC^-CITRINE expressed under the *SHR* promoter showed no epidermal signal (Fig. S8C) and conferred no rescue (Fig. S8D), but higher levels triggered excess formative divisions in the stele (Fig. 5H, Fig. S8E) consistent with previous reports (Fridman et al., 2021, Graeff et al., 2021, Kang et al., 2017). Such excess divisions were not observed when BRI1^REC^-CITRINE was expressed under the control of the other promoters, suggesting that they reflect local effects of (hyperactive) brassinosteroid signaling. Finally, we also expressed BRI1^REC^-CITRINE under the control of the epidermis-specific *ARABIDOPSIS THALIANA MERISTEM LAYER 1 (ATML1)* promoter (Sessions et al., 1999) (Fig. S1), which had been used in the original transgenic complementation of *bri1* shoot growth (Savaldi-Goldstein et al., 2007). However, unlike in *ATML1::BRI1-CITRINE* controls, neither shoot nor root growth defects were complemented in *ATML1::BRI1*^*REC*^*-CITRINE* lines (Fig. 5A and I) despite epidermal BRI1^REC^-CITRINE signal (Fig. 5J-K). Moreover, the BRI1-CITRINE signal was spread more evenly throughout the epidermis whereas the BRI1^REC^-CITRINE signal resembled the *ATML1* expression pattern (Fig. 5L, Fig. S1). Together, these results indicate that expression of the brassinosteroid receptor in the *ATML1* expression domain of the epidermis is not by itself sufficient for *bri*^*3*^ rescue.

## DISCUSSION

Loss-of-function genetics has established BRI1 as the dominant Arabidopsis brassinosteroid receptor because it is essential for growth (Li and Chory, 1997, Li et al., 2022), whereas minor roles or specific functions were assigned to its homologs BRL1 and BRL3 (Cano-Delgado et al., 2004, Fabregas et al., 2018). In *bri1* single mutants, *BRL1* and *BRL3* are presumably still active in stele tissues. Rescue of the *bri*^*3*^ phenotype by BRL1 or BRL3 expression under control of tissue-specific promoters may therefore appear counterintuitive. However, we have shown previously that the extent of *bri*^*3*^ rescue through phloem-specific *BRI1* expression depends on expression levels and transgene copy number (Graeff et al., 2020, Graeff et al., 2021), and even single locus transgene insertions are typically concatenated (Dickinson et al., 2023). Thus, our data reiterate the importance of quantitative brassinosteroid signaling and indicate that *BRL1* and *BRL3* expression is normally below the threshold to compensate for the absence of *BRI1*. Moreover, differences in rescue efficiency may also reflect the local impact of brassinosteroid signaling on brassinosteroid homeostasis. For example, the rescue observed with phloem pole-specific promoters may be enhanced by the regulation of genes encoding rate-limiting brassinosteroid biosynthesis enzymes, which are expressed in the vascular cylinder (Vukasinovic et al., 2021, Ohashi-Ito et al., 2023).

A central deduction from our combination of promoters and *BRI1* coding sequences in genetic analyses (Fig. 5M) is that DNA sequences in the coding region of brassinosteroid receptor genes contribute to their expression pattern and confer low expression throughout root tissues. This is most evident in the ectopic receptor signal observed with stronger phloem-specific promoters, which also suggests a synergism between regulatory sequences in the promoter and the gene body. The contribution of the transcript region to gene expression pattern has so far only been observed rarely (Cattaneo et al., 2019, Koh et al., 2021, Sieburth and Meyerowitz, 1997). A comprehensive comparison of promoter-driven reporter gene expression patterns with scRNA-seq data could reveal how common this phenomenon is.

Collectively, our confocal microscopy, scRNAseq and genetic data suggest that *BRI1* transgenes are weakly expressed throughout root tissues and that trace amounts of BRI1 are frequently sufficient to sustain root growth in the *bri*^*3*^ background. This is surprising given that the ectopic BRI1-CITRINE/GFP signal can be below the detection threshold of contemporary confocal microscopes. Generally, it was only clearly evident as above-background fluorescence plasma-membrane-localized signal in the absence of any counterstaining. The necessity of the ectopic epidermal expression for *bri*^*3*^ complementation reiterates the importance of the epidermis in restricting organ growth (Savaldi-Goldstein et al., 2007). Yet our results also suggest that a threshold level of brassinosteroid perception across multiple, if not all tissues is necessary for comprehensive *bri*^*3*^ rescue. The results may also mean that a lower level of brassinosteroid signaling across tissues is preferable over a strong imbalance between inner and outer cell layers, which may sometimes prevent phenotypic recovery as previously suggested (Fridman et al., 2021, Vragovic et al., 2015). Conversely, restricted expression of the brassinosteroid receptor in individual tissues is not sufficient to normalize the *bri*^*3*^ mutant phenotype. The most parsimonious interpretation of our data is therefore that brassinosteroid receptors largely act in a cell-autonomous manner.

## ACKNOWLEDGEMENTS

This study was supported by Swiss National Science Foundation grant 310030L_19779 awarded to C.S.H. and Research Foundation-Flanders Postdoctoral fellowships 12R7822N and 12R7819N awarded to N.V. T.M.N. and C.W.H. acknowledge support from the Howard Hughes Medical Institute, where Prof. Philip Benfey was an investigator. We honor the memory of Philip Benfey, whose contributions laid the groundwork for this research.

## MATERIALS AND METHODS

### Plant materials and growth conditions

All the lines used in this work were in the Arabidopsis wildtype accession Columbia-0 (Col-0) background. The *bri1-116 brl1 brl3* triple mutant (*bri*^*3*^) and the *CVP2::BRI1-CITRINE bri*^*3*^, *BAM3::BRI1-CITRINE bri*^*3*^, *MAKR5::BRI1-CITRINE bri*^*3*^, *SHR::BRI1-GFP bri*^*3*^, *GL2::BRI1-GFP bri*^*3*^ and *BRI1::BRI1-GFP bri*^*3*^ transgenic lines were described previously (Graeff et al., 2020, Kang et al., 2017, Vragovic et al., 2015). For *in vitro* culture, seeds were surface sterilized, then stratified at 4°C for 2-3 days. Seeds were placed on plates containing ½ Murashige and Skoog medium (MS, including vitamins and MES buffer; Duchefa, M0255) supplemented with 1% agar and 0.3% sucrose. The pH was adjusted to 5.7. Seedlings were grown vertically in a growth chamber under continuous white light of c. 120 µmol m^-2^ s^-1^ intensity at 22°C. CLE peptides were obtained from a commercial supplier (Genscript; synthesized at >80% purity), diluted in water and added to the medium at a final concentration of 15nM. For root growth assays, brassinolide (Sigma Aldrich, Product No. SML0094) and brassinazole (Tokyo Chemical Industry, Product No. B2829) were applied at the indicated final concentrations. For root growth measurements, plates scanned with a high-resolution flatbed scanner were analyzed using the Simple Neurite Trace plug-in for Fiji software. All the experiments were repeated at least once.

### Transgenic brassinosteroid receptor and reporter lines

For transgene constructs, attB-flanked coding sequences were synthetized (*BRI1*^*REC*^, *BRI1*^*KDD*^) or amplified (*BIR3*^*EXT*^*-BRI1*^*INT*^, *BRI1, BRL1, BRL2* and *BRL3*) by PCR and recombined into the pDONR221 vector (Invitrogen) to produce the pEN-L1-CDS-L2 clones. Protein fusions under the control of different promoters were generated by recombining pEN-L4-promoter-R1, pEN-L1-CDS-L2 and pEN-R2-CITRINE-L3 plasmids into a destination vector by Multisite Gateway LR reaction (Thermo Fisher Scientific). The *BRI1, CLE45, COR, CVP2, ATML1* and *SHR* promoters have been previously described (Gallego-Bartolome et al., 2011, Kang et al., 2017, Nolan et al., 2023, Savaldi-Goldstein et al., 2007, Sessions et al., 1999, Vragovic et al., 2015). The destination vectors used were pFR7m34GW (for *CLE45::NLS-SCARLET* and *BRI1*^*REC*^ constructs), pK8m34GW-FAST (for *COR::BRI1-GFP*) and pH7m34GW (for all other constructs). All constructs were verified by Sanger sequencing and then introduced into *Agrobacterium tumefaciens* strain GV3101 with the pMP90 helper plasmid. In order to obtain the different transgenes in *bri*^*3*^ background, homozygous *brl1 brl3* double mutant plants that were heterozygous for the *bri1-116* allele were transformed by the floral dip method. Transgenics in homozygous *bri*^*3*^ background were selected by genotyping as described (Graeff et al., 2020, Vragovic et al., 2015). The *COR::BRI1-GFP* construct was transformed into a *WOX5::erGFP bri*^*3*^ background, the *CLE45::NLS-SCARLET* construct also into Col-0. For each construct-background combination several independent transgenic lines were obtained. Observations were typically confirmed in detail in two or more independent transgenic lines.

### BRI1 transgene knockout by tissue-specific CRISPR/Cas9

The *pEN-R2-gRNA_BRI1-3-gRNA_BRI1-2-L3* vector, which contains two gRNAs targeting *BRI1* as well as the *WER::Cas9*^*BRI1*^ expression construct have been previously described (Nolan et al., 2023). The *SHR::Cas9*^*BRI1*^ expression clone was generated by combining the pDONRL4-L1r plasmid carrying the *SHR* promoter, the *pEN-R2-gRNA_BRI1-3-gRNA_BRI1-2-L3* plasmid, and the destination vector pK8m34GW-FAST in a MultiSite Gateway LR reaction. *pCVP2::BRI1-CIT bri*^*3*^, *pCLE45::BRI1-CIT bri*^*3*^ and *pMAKR5::BRI1-CIT bri*^*3*^ plants were transformed with both *WER::Cas9*^*BRI1*^ and *SHR::Cas9*^*BRI1*^ expression constructs by the floral dip method. Transgenic T2 generation plants were selected based on the presence of GFP signal in the seed coat, and the efficiency of the tissue-specific CRISPR was confirmed by confocal microscopy.

### DWF1 knockout by CRISPR/Cas9

Three gRNAs (5’-ACT CTG ACC ACA TGT CCC CG-3’, 5’-ATC TTT ACT ACG CAA TCC CG-3’, 5’-CTA CTT CCT CAT CTA CCT CG-3’) were designed to target the first exon of *DWF1* (AT3G19820) in the *pCVP2::BRI1-CITRINE bri*^*3*^ line. T1 generation plants were selected based on FASTRED, and the mutations were confirmed by genotyping and sequencing. The following primers were used: *2F*, 5’-TGG TTT GAT GCA GTG A-3’, *DWF1 crispr geno 2R*, 5’-CAC GGC TTG AAC CAC-3’. Different alleles were found, and the hypomorhpic *dwf1*^*del48*^ allele was chosen for further experiments because it is able to produce seeds.

### Confocal microscopy

Live confocal microscopy was performed in most cases. For detection of fluorescent proteins, the following emission-excitation wavelengths in a *Leica Stellaris 5* instrument were used: excitation 488 or 514nm / emission 493–565nm (GFP/CITRINE), and excitation 561nm / emission 566–734nm (NLS-SCARLET). Formative cell divisions were analyzed in 7-day-old roots fixed with 4% paraformaldehyde in PBS buffer for 30min and cleared with *ClearSee* solution for five days. Cleared roots were stained with 0.1% Calcofluor White (CAS-No: 4193-55-9; Sigma-Aldrich) in *ClearSee* solution and washed two times with PBS buffer (10-15min ea.). Samples were imaged on a *Leica Stellaris 5* confocal microscope with 20x and 63x objectives, using the 405nm laser for calcofluor excitation. For image analyses, Fiji software was used.

### 10X Genomics scRNA-seq of Arabidopsis root protoplasts

scRNAseq analysis of *CVP2::BRI1-CITRINE* in *bri*^*3*^ was performed in a side-by-side experiment along with published samples for wild-type, *bri*^*3*^, and *GL2::BRI1-GFP* in *bri*^*3*^ as previously described (Nolan et al., 2023). Plants were grown vertically in a growth chamber set to 22°C, 16 hours light/8 hours dark for 7 days on 1/2 Linsmaier and Skoog (LSP03-1LT, Caisson Labs; pH 5.7) 1% sucrose media with 100μm nylon mesh (Nitex 03-100/44). Root tips were harvested from 1000-3000 roots per sample by cutting c. 0.5cm from the tip with a razor blade. Excised roots were placed into a 35mm petri dish containing a 70μm cell strainer and 4.5mL enzyme solution (1.5% [w/v] cellulase [ONOZUKA R-10, GoldBio], 0.1% Pectolyase [Sigma P3026], 0.4M mannitol, 20mM MES (pH 5.7), 20mM KCl, 10mM CaCl2, 0.1% bovine serum albumin, and 0.000194% (v/v) beta-mercaptoethanol). The digestion was incubated on an 85-rpm shaker at 25°C for one hour with additional stirring every 15-20 minutes. The resulting cell solution was filtered twice through 40μm cell strainers and centrifuged for 5 minutes at 500g in a swinging bucket rotor. The pellet was washed with 2mL washing solution (0.4M mannitol, 20mM MES (pH 5.7), 20mM KCl, 10mM CaCl2, 0.1% bovine serum albumin, and 0.000194% (v/v) beta-mercaptoethanol), centrifuged again at 500g for 3 minutes, and the pellet resuspended in washing solution at a concentration of c. 2000 cells/µL. We loaded 16,000 cells, with the aim to capture 10,000 cells per sample with the *10X Genomics* Chromium 3’ Gene expression v3.1 kits. Cell barcoding and library construction were performed following the manufacturer’s instructions. cDNA and final library quality were verified using a *Bioanalyzer* High Sensitivity DNA Chip (Agilent) and sequenced on an *Illumina NovaSeq 6000* instrument.

### scRNAseq data processing and analysis

scRNAseq analysis was carried out as described (Nolan et al., 2023), except that the genome sequences were modified to analyze the *BRI1* transgenes. Sequencing reads were demultiplexed from *Illumina* BCL files to produce FASTQ files for each sample using *CellRanger* mkfastq (v3.1.0, *10X Genomics*). We then created two separate custom reference files using the Arabidopsis TAIR10 reference genome. The first contained the *BRI1-CITRINE* transgene sequence, which was used to analyze wild type and *CVP2::BRI1-CITRINE* in *bri*^*3*^, while the second contained *BRI1-GFP* and was used to analyze wild type and *GL2::BRI1-GFP* in *bri*^*3*^. Reads were then aligned against the custom reference to generate a gene-by-cell matrix using the scKB script (https://github.com/ohlerlab/scKB), which incorporates *kallisto* and *bustools* (Bray et al., 2016, Melsted et al., 2019). Quality filtering of cells was performed using the R package *COPILOT* (Cell preprOcessing PIpeline kaLlistO busTools) (Hsu et al., 2022), which uses a non-arbitrary scheme to remove empty droplets and dying or low-quality cells. One iteration of *COPILOT* filtering adequately separated high-quality cells from the background in these samples based on an examination of barcode rank plots. The resulting high-quality cells were further filtered to remove outliers based on the top 1% of cells in terms of UMI counts, and putative doublets were removed with *DoubletFinder* (McGinnis et al., 2019), incorporating the estimated doublet rate from the *10X Genomics* Chromium Single Cell 3’ Reagent Kit user guide. In total, we identified 20,957 high-quality cells from two biological replicates of *CVP2::BRI1-CITRINE* in *bri*^*3*^.

Normalization, annotation, and integration of scRNAseq datasets were carried out using *Seurat*. Data were normalized using *SCTransform* (Hafemeister and Satija, 2019). All genes except those from mitochondria, chloroplasts, or those affected by protoplasting (Denyer et al., 2019, Shahan et al., 2022) (absolute log2 fold-change >=2) were retained for analysis. Cell type and developmental stage labels from the wild-type atlas (Nolan et al., 2023, Shahan et al., 2022) were transferred to each sample via label transfer in *Seurat* (Butler et al., 2018, Stuart et al., 2019). We integrated the samples from each custom reference using the *Seurat* integration pipeline. A sample from the atlas with the highest number of detected genes (sc_12) (Shahan et al., 2022) and two previously described samples (dc_1 and dc_2) (Denyer et al., 2019) were included in the integration to facilitate comparable visualizations but were excluded from any downstream analysis. Principal component analysis (PCA) was performed by calculating 50 principal components using the RunPCA function (with approx=FALSE). UMAP non-linear dimensionality reduction was next calculated via the RunUMAP function using all 50 principal components with parameters n_neighbors = 30, min_dist = 0.3, umap.method = ‘‘umap-learn’’, metric =‘‘correlation’’ using the “integrated” assay. These processing steps have been previously described (Nolan et al., 2023, Shahan et al., 2022) and are documented in jupyter notebooks as part of the COPILOT workflow. Gene expression patterns were examined by plotting the normalized expression values produced by the SCTransform function, with the 90^th^ quantile of expression as the maximum cutoff. To quantify *BRI1* transgene levels in different cell types, we used *muscat* (multi-sample multi-group scRNA-seq analysis tools) (Crowell et al., 2020) to aggregate cell level counts for each cell type on a per-sample basis. Raw counts were summed using the aggregateData function and differential expression testing was performed using *edgeR* (McCarthy et al., 2012) incorporated in the pbDS function of *muscat*. A gene was considered differentially expressed in a given cell type if the false discovery-rate adjusted p-value was <=0.05, absolute fold change was >=1.5 and detection frequency was >=5% in one of the genotypes. Tables were exported from muscat with counts per million normalized expression values for each cell type/sample combination.

## Statistical analysis

Data was analyzed using GraphPad Prism software version 10.2.1. Robust regression and Outlier removal (ROUT) analyses were performed on root measurements to detect (rare) outliers, which were removed. Specific statistical tests used are indicated in the figure legends and were always two-sided.

## Data and Code Availability

The published article includes all datasets generated or analyzed during this study, except the raw and processed scRNA-seq data, which have been deposited at the NCBI GEO under accession GSE212230. The code to reproduce this study can be found at https://github.com/tmnolan/BR-Receptors. These data will be made publicly accessible in the Arabidopsis Root Virtual Expression eXplorer (ARVEX; https://shiny.mdc-berlin.de/ARVEX/).

---

**Fig. S1.**
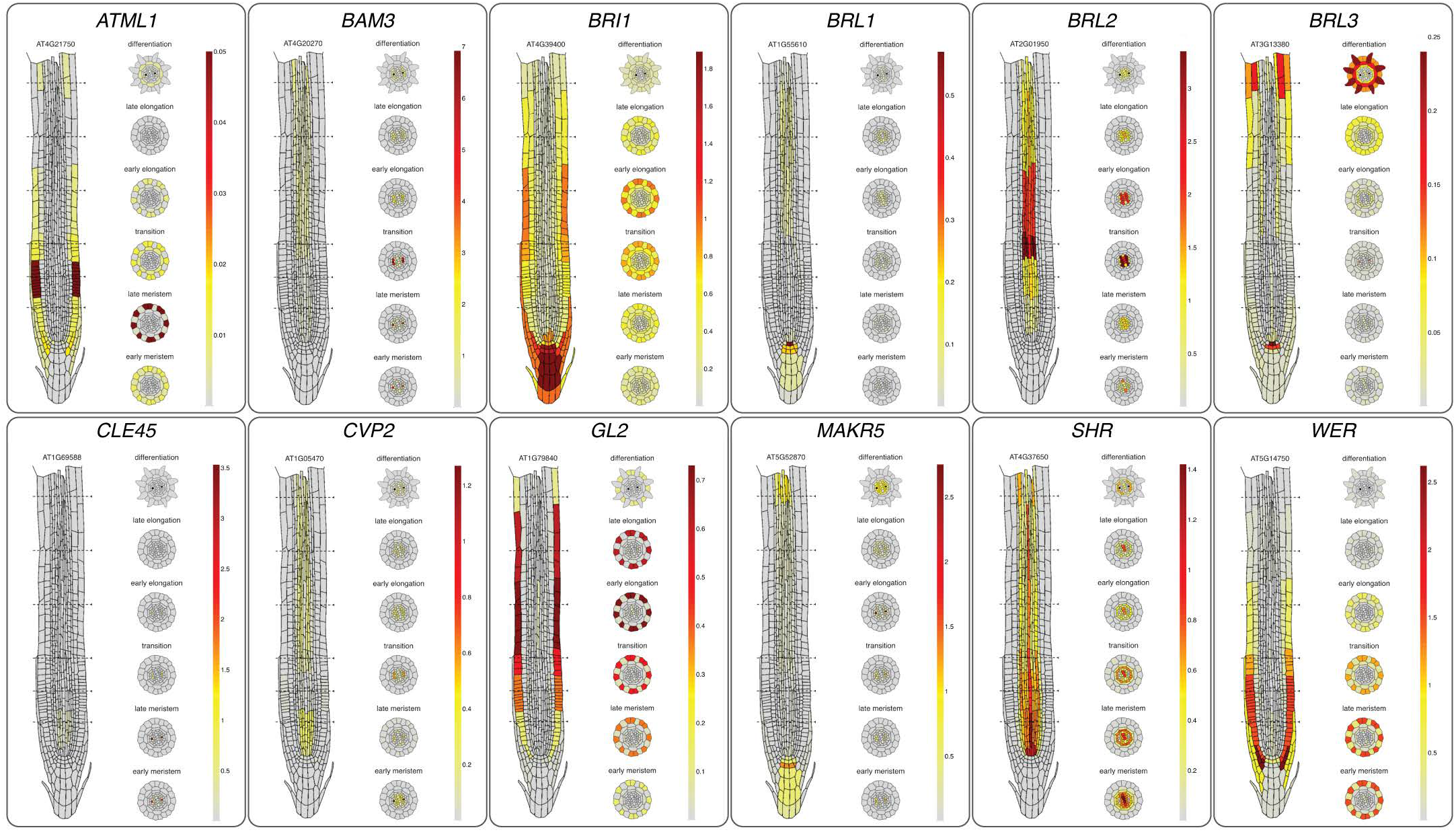
Expression patterns of genes investigated in this study. Schematic representation of root tip expression patterns of indicated genes, obtained from aggregation of multiple independent single cell mRNA analyses of Arabidopsis Col-0 wildtype roots (https://rootcellatlas.org). Note the differences in expression level scales.

**Fig. S2.**
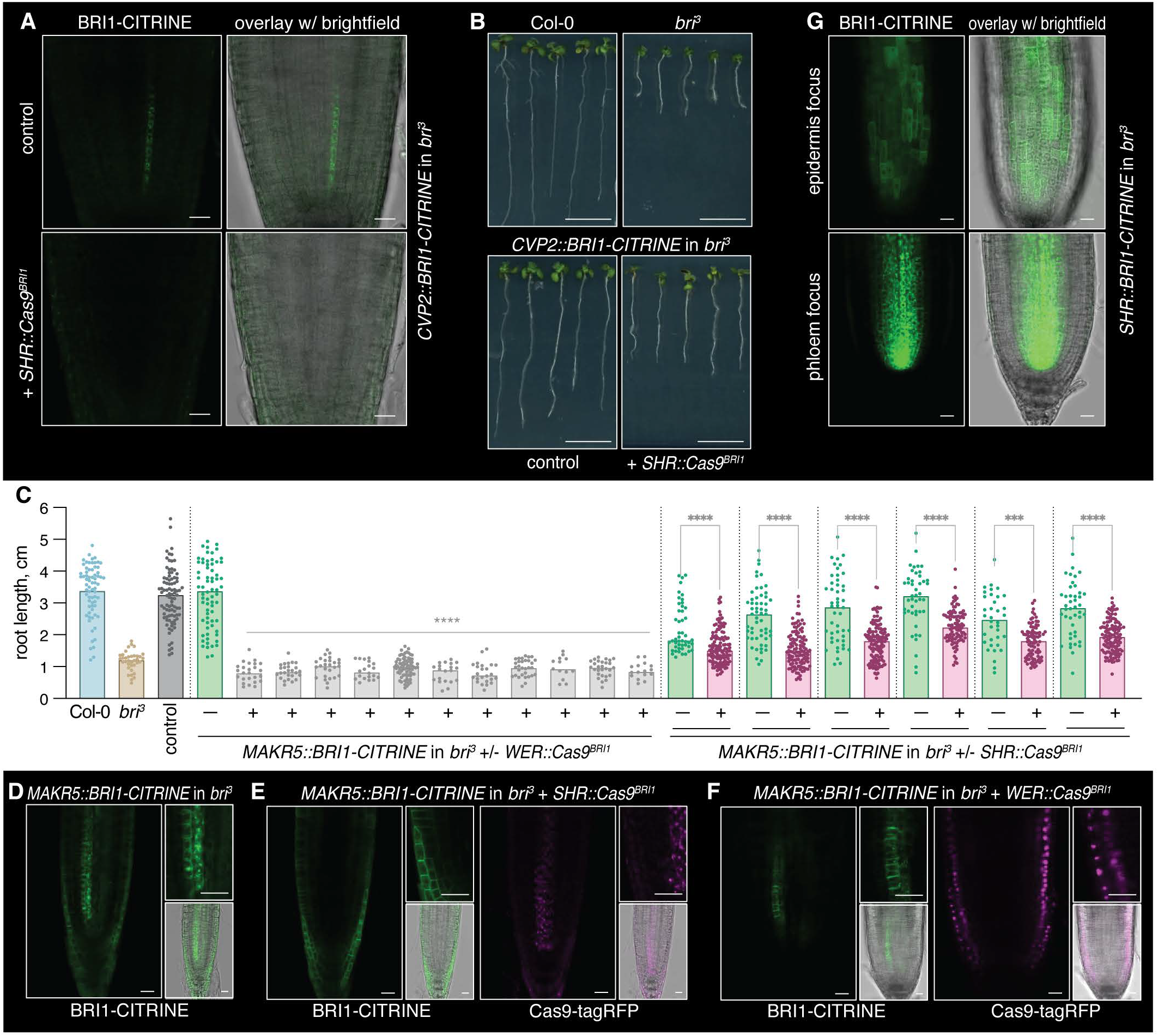
Trace amounts of epidermal BRl1 expression are required for *bri*^*3*^ rescue. **(A)** Top: Confocal microscopy images of a root meristem from a *bri*^*3*^ seedling complemented with a BRl1-CITRINE fusion protein (green fluorescence) expressed under control of the phloem sieve element-specific *CVP2* promoter, imaged with relaxed settings and without any counterstaining. Bottom: CRISPR/Cas9 knockout of the *BRl1* transgene using the stele-specific *SHR* promoter leads to disappearance of BRl1-CITRINE signal. **(B)** Representative 8-day-old *bri*^*3*^ seedlings complemented with a *CVP2::BRl1-CITRINE* transgene (bottom left) and the same line combined with stele-specific CRISPR/Cas9 *BRl1* knockout (bottom right) as compared to controls (top). **(C)** Root growth quantification for 8-d-old seedlings of the indicated genotypes. A *bri*^*3*^ line complemented with *BRl1-CITRINE* expressed under control of the phloem pole-specific *MAKR5* promoter (control) was combined with tissue-specific CRISPR/Cas9 *BRl1* knockout using either the stele-specific *SHR* or the epidermis-specific *WER* promoter, several independent lines are shown. Statistically significant differences (asterisk) between seedlings carrying the CRISPR/Cas9 construct and their segregating non-transgenic siblings were determined by ordinary one-way ANOVA, p<0.0001. **(D-F)** Confocal microscopy images of root meristems from *bri*^*3*^ seedlings complemented with *BRl1-CITRINE* expressed under control of the *MAKR5* promoter (D) and combined with tissue-specific CRISPR/Cas9 *BRl1* knockout using the *SHR* (E) or *WER* (F) promoters, showing both BRl1-CITRINE and Cas9-tagRFP signals. **(G)** Confocal microscopy images of a root meristem from a *bri*^*3*^ seedling expressing BRl1-CITRINE fusion protein under control of the *SHR* promoter. Size bars are 20 µ m in microscopy images and 1cm in seedling images.

**Fig. S3.**
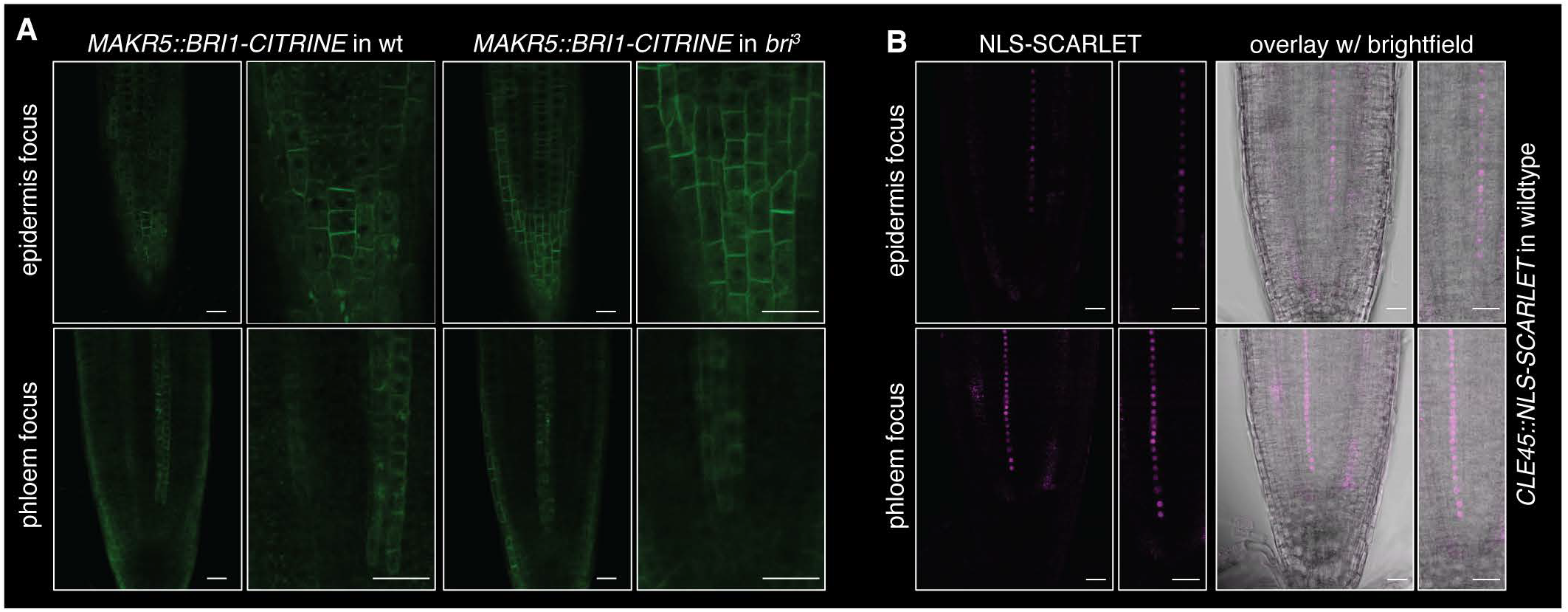
Trace epidermal BRl1 expression is independent of the promoter. **(A)** Comparison of phloem pole-specific *MAKR5* promoter-driven **BRl1-** CITRINE signal in morphologically wildtype *(bri1* +/− *brl1* +/− *brl3* ***+/−)*** and *bri*^*3*^ background. **(B)** Confocal microscopy of an NLS-SCARLET fusion protein expressed under control of the phloem sieve element-specific *CLE45* promoter in Col-0 wildtype background. Size bars are 20 µ m.

**Fig. S4.**
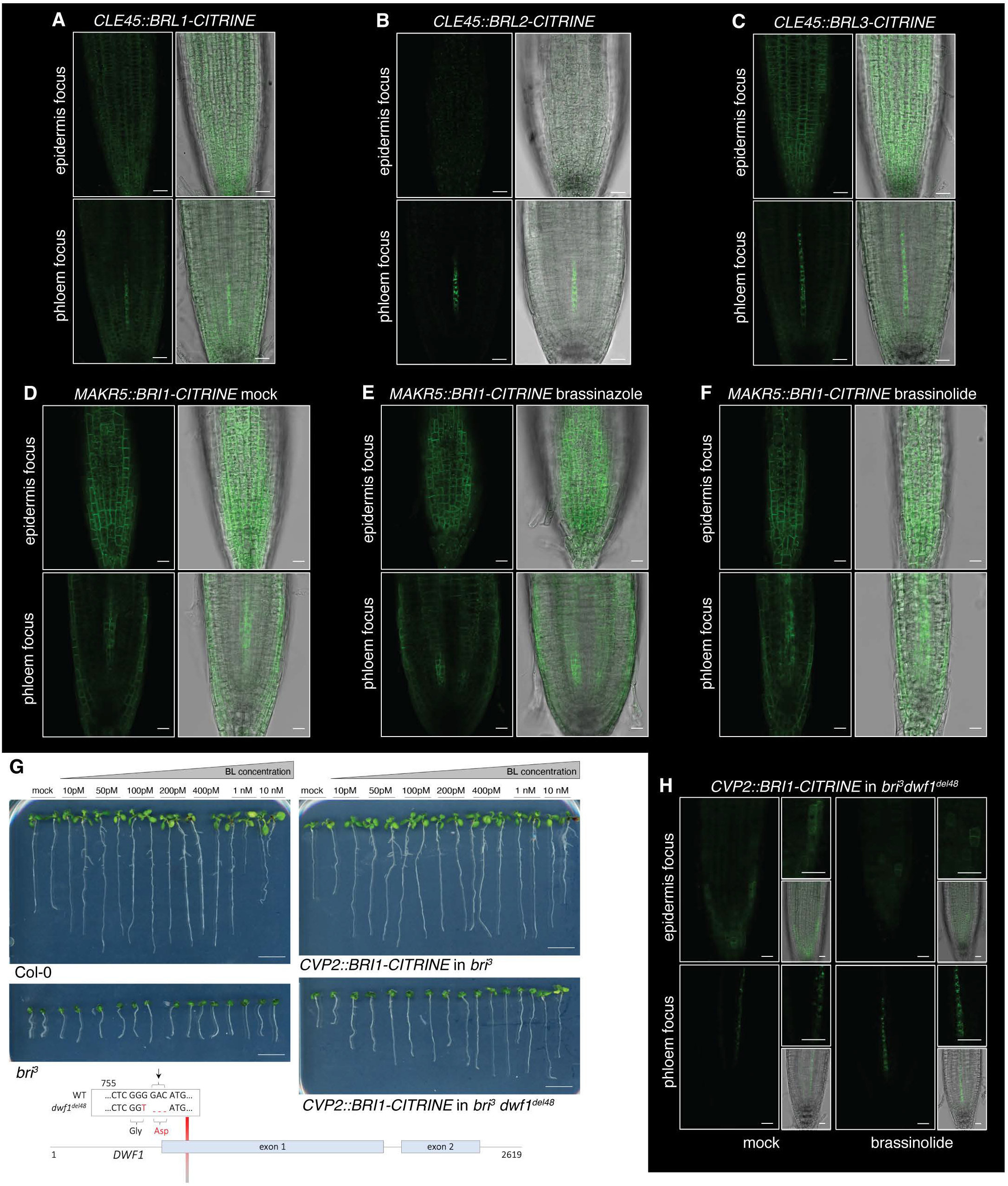
Trace epidermal BRl1 expression does not result from autoregulatory feedback. **(A-C)** Confocal microscopy images of root meristems from *bri*^*3*^ seedlings expressing CITRINE fusion proteins (green fluorescence) of the indicated BRl1 homologs under control of the phloem sieve element-specific *CLE45* promoter. **(D-F)** Confocal microscopy images of root meristems from *bri*^***3***^ seedlings expressing BRl1-CITRINE fusion protein under control of the phloem pole-specific *MAKRS* promoter, treated with mock, 1mM brassinazole, or 10nM brassinolide. **(G)** Representative 9-day-old *bri*^*3*^ seedlings complemented with a *CVP2::BRl1-CITRINEtransgene* and the same line combined with a hypomorphic mutation in the brassinosteroid biosynthesis gene *DWARF1 (dwf1*^*del48*^; see schematic), compared to controls and treated with increasing levels of brassinolide. **(H)** Confocal microscopy images of a root meristem from *bri*^*3*^ *dwft*^*del48*^ quadruple mutant seedlings carrying a *CVP2::BRl1-CITRINE* transgene, treated with mock or 100nM brassinolide. Size bars are 20 µ m in microscopy images and 1cm in seedling images.

**Fig. S5.**
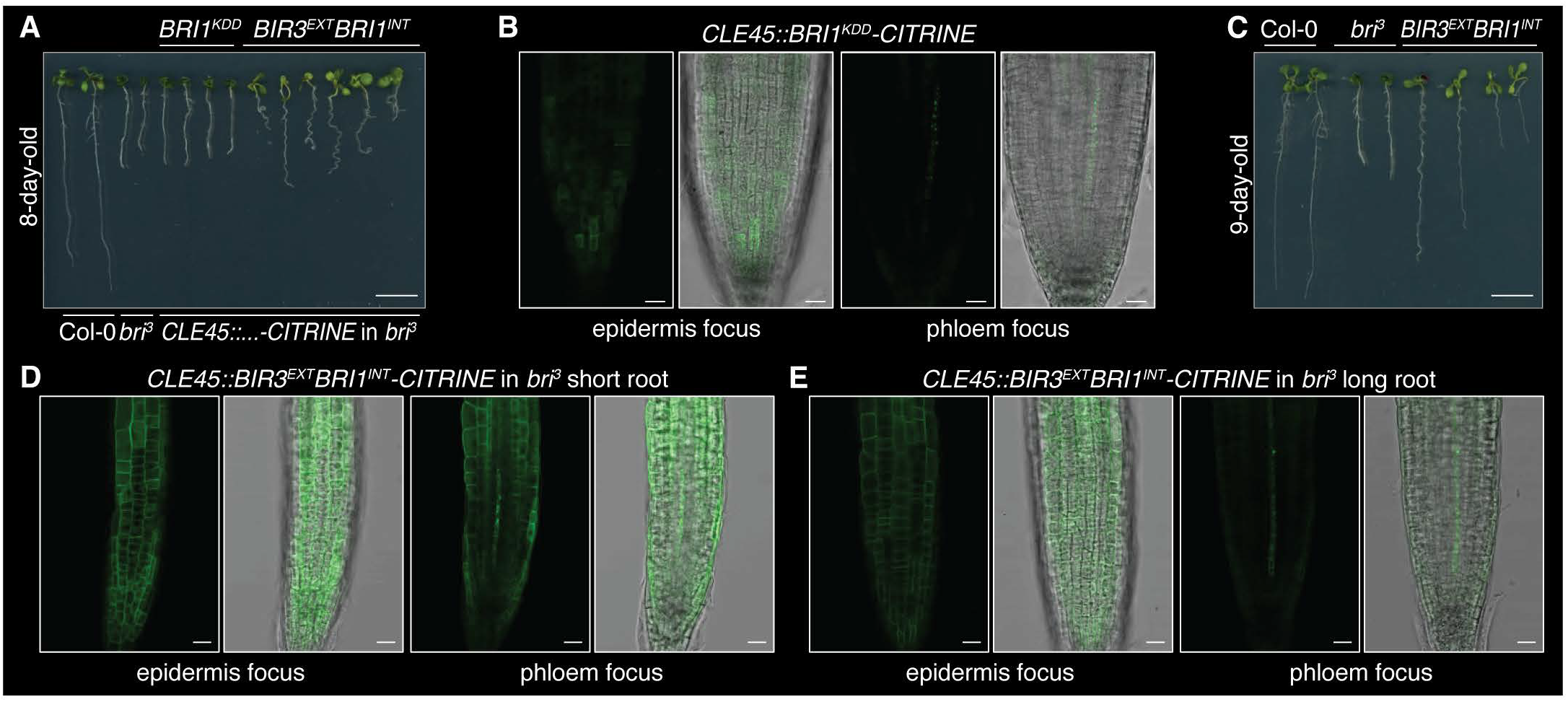
Active BRl1 kinase domain is necessary and sufficient for *bri*^*3*^ complementation. **(A)** Representative 8-day-old brf3 seedlings expressing CITRINE fusion proteins with an inactive (BRl1 ^KDD^-CITRINE) or a constitutively active (BIR3^EXT^BRl1^INT^) BRl1 kinase domain under control of the phloem sieve element-specific CLE45 promoter, compared to controls. **(B)** Confocal microscopy images of a root meristem from a brf^3^ mutant seedling carrying a CLE45::BRJ1^KDD^-C/TRINE transgene. **(C)** Representative 9-day-old brf^3^ seedlings carrying a CLE45::BIR3^EXT^BRl1^*INT*^-CJTRINE transgene, compared to controls, illustrating the phenotypic range observed (also see (A)). **(D-E)** Confocal microscopy images of root meristems from brf3 mutants expressing the CLE45::BIR3^EXT^BRl1^*INT*^-C/TRINE transgene across the phenotypic range illustrated in (C). Note the ubiquitous fusion protein signal. Size bars are 20 µ m in microscopy images and 1cm in seedling images.

**Fig. S6.**
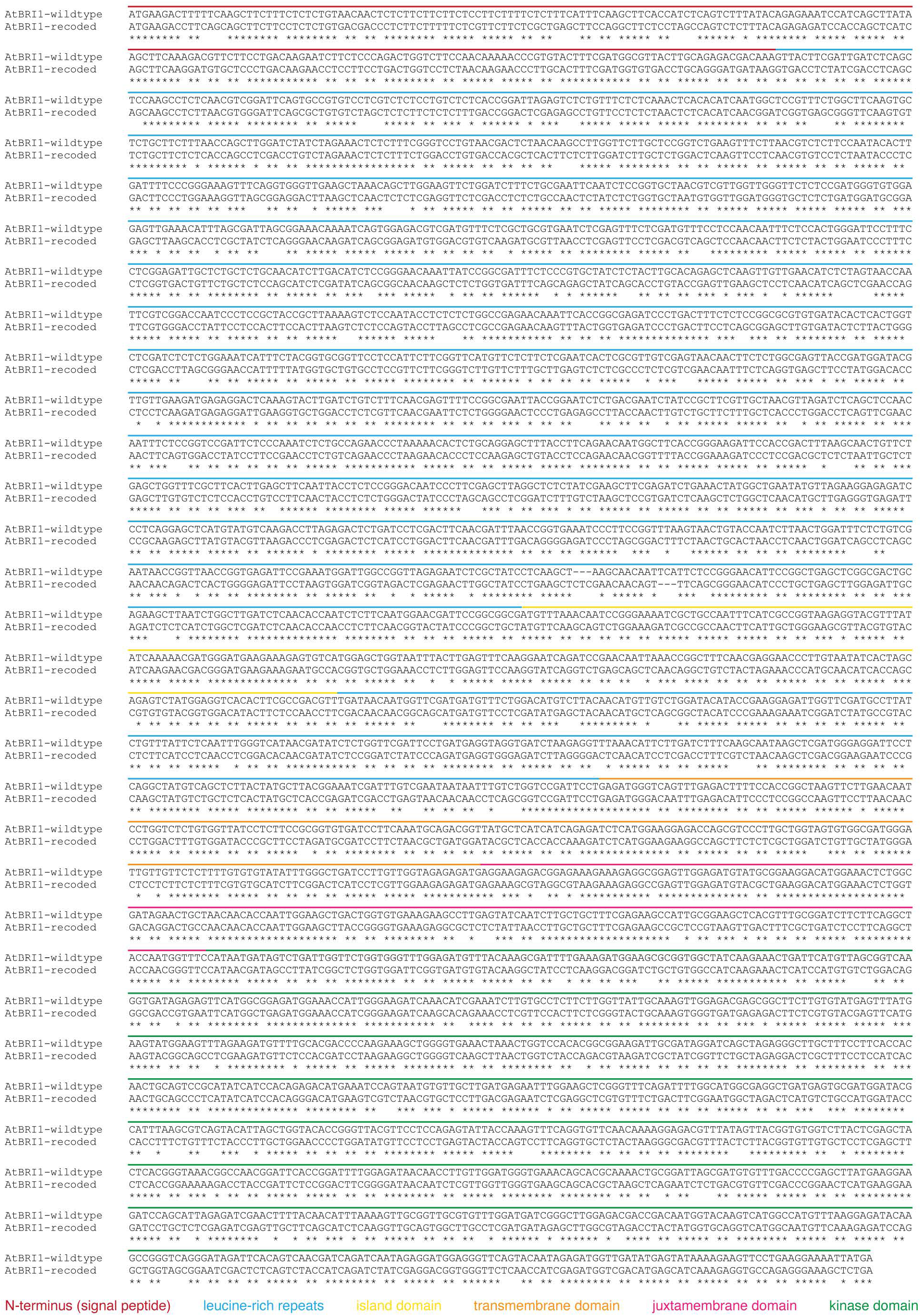
Recoded *BRl1* gene body sequence. Alignment of the native *BRl1* coding sequence and the recoded version, with the corresponding protein domains indicated. Note that the coding sequence of *BRl1* and other brassinosteroid receptor genes is not interrupted by any intrans.

**Fig. S7.**
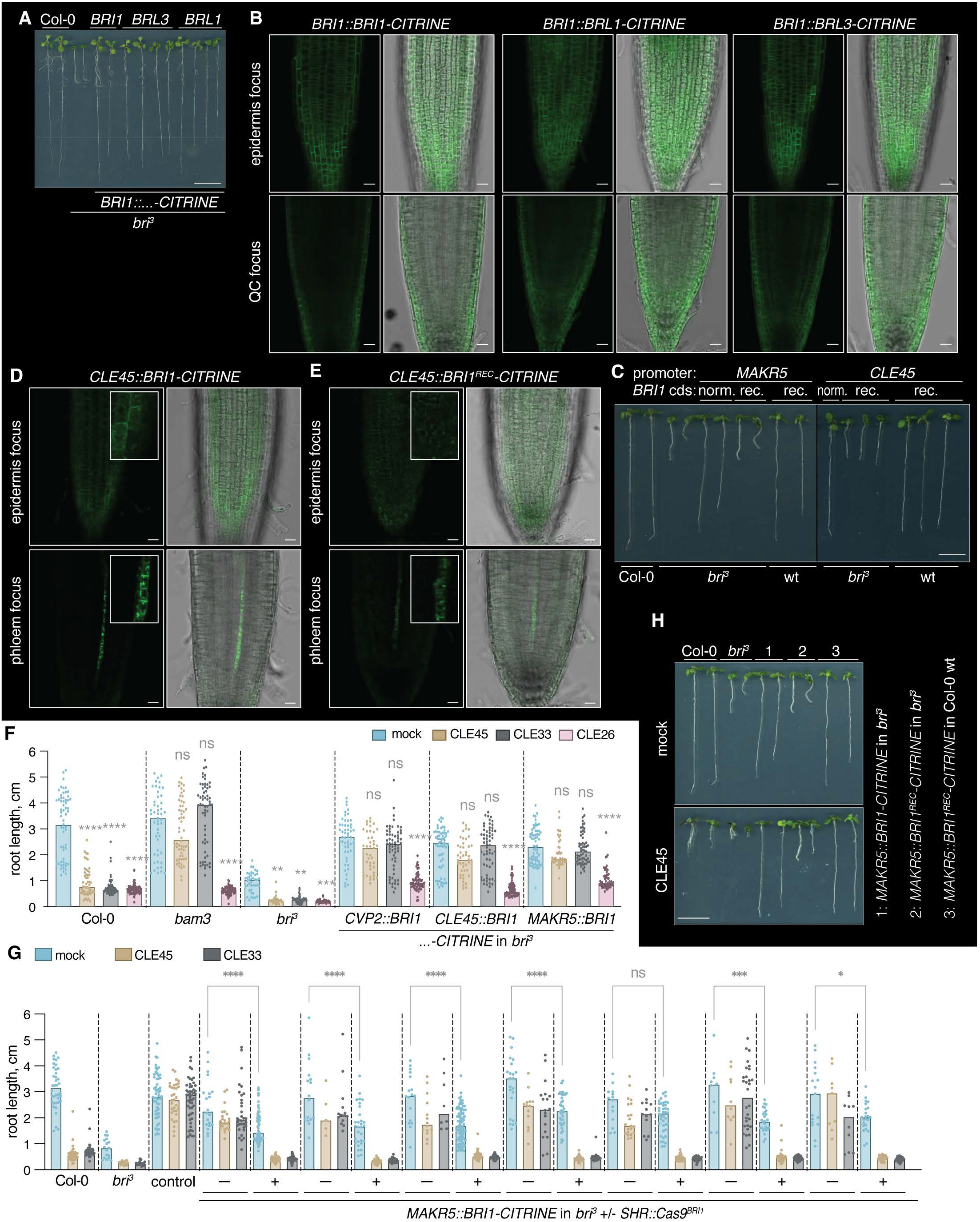
Recoded *BRl1* transgenes produce functional BRl1 fusion protein but cannot reproduce *bri*^*3*^ complementation when expressed under control of tissue-specific promoters. **(A)** Representative 9-d-old *bri*^*3*^ seedlings expressing brassinosteroid receptors under control of the native *BRl1* promoter, compared to wildtype and *bri*^*3*^ controls. **(B)** Confocal microscopy images of root meristems from seedlings shown in (A). **(C)** Representative 8-d-old *bri*^*3*^ or morphologically wildtype *brl1 brl3* double mutant seedlings carrying different *BRl1-CITRINEtransgenes* made with the genuine (norm.) or recoded (rec.) *BRl1* coding sequence (eds), compared to wildtype and *bri*^*3*^ controls. **(D-E)** Confocal microscopy images of root meristems from seedlings expressing BRl1-CITRINE or BRl1^REC^-cITRINE fusion protein under control of the *CLE45* promoter in *bri* mutants, illustrating the absence of epidermal BRI1 ^REC^ –cITRINE signal. **(F-G)** Root growth quantification for 7-d-old seedlings of the indicated genotypes grown on mock or in the presence of 20nM of the indicated CLE peptides as compared to controls. (F) The *bam3* mutant is specifically insensitive to CLE45 and the control peptide CLE33, but sensitive to the control peptide CLE26. *bri*^*3*^ lines complemented with *BRl1-CITRINE*expressed under control of phloem (pole)-specific promoters behave similarly. (G) A *bri*^*3*^ line complemented with *BRl1-CITRINE*expressed under control of the phloem pole-specific *MAKR5* promoter (control) was combined with tissue-specific CRISPR/Cas9 *BRl1* knockout using the stele-specific *SHR* promoter, several independent lines are shown. Statistically significant differences (asterisks) between seedlings carrying the CRISPR/Cas9 construct and their segregating non-transgenic siblings were determined by ordinary one-way ANOVA, JX0.001. **(H)** Representative 9-d-old mock (top) or CLE45 peptide-treated (20nM) (bottom) *bri*^*3*^ or morphologically wildtype *brl1 brl3* double mutant seedlings expressing BRl1-CITRINE or BRl1 ^REC^_CITRINE fusion protein under control of the *MAKR5* promoter, compared to controls. Size bars are 20 µm in microscopy images and 1cm in seedling images.

**Fig. S8.**
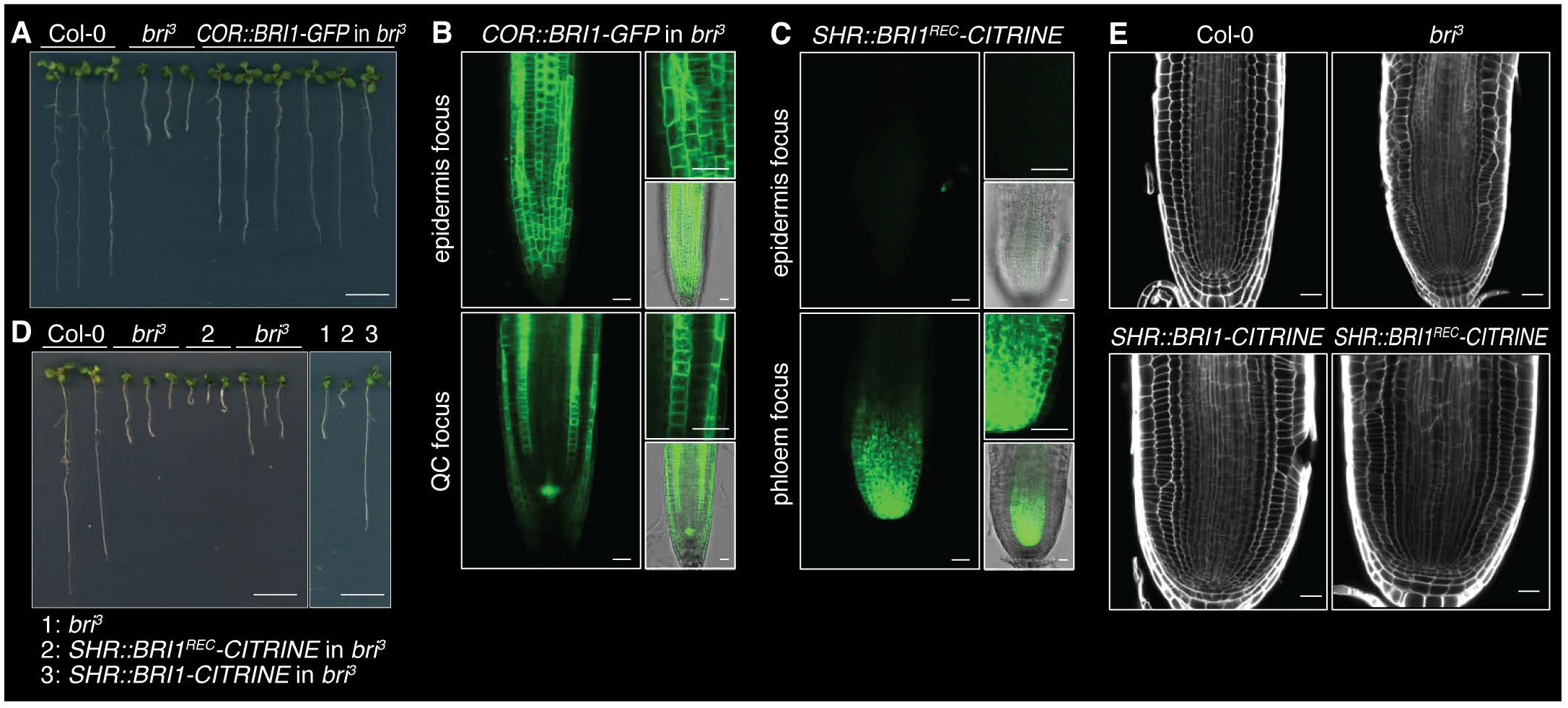
Phloem-specific increase of *BRl1* activity confers resistance to CLE45 peptide application. **(A)** Representative 8-day-old *bri*^*3*^ seedlings complemented with a BRl1-GFP fusion protein expressed under control of the cortex-specific CO promoter, compared to controls. **(B)** Confocal microscopy images of a root meristem from seedlings shown in (A). Note the presence of a background marker gene expressed in the quiescent center (QC). **(C)** Confocal microscopy images of a root meristem from a *bri*^*3*^ mutant expressing BRl1^REC^-CITRINE fusion protein under control of the stele-specific *SHR* promoter. **(D)** Representative 8-day-old seedlings expressing BRl1-CITRINE or BRl1 ^REC.^-CITRINE fusion protein under control of the *SHR* promoter in *bri*^*3*^ mutants, compared to controls. **(E)** Confocal microscopy images of calcofluor-stained root meristems from *bri*^*3*^ mutants and their counterparts expressing either BRl1-CITRINE or BRl1^REC.^-CITRINE fusion protein under control of the *SHR* promoter. Right panels reproduced from Figure 5. Size bars are 20 µ m in microscopy images and 1cm in seedling images.

## REFERENCES

Bray, N. L., Pimentel, H., Melsted, P. & Pachter, L. 2016. Near-optimal probabilistic RNA-seq quantification. Nat Biotechnol, 34, 525–7.

Breda, A. S., Hazak, O., Schultz, P., Anne, P., Graeff, M., Simon, R. & Hardtke, C. S. 2019. A Cellular Insulator against CLE45 Peptide Signaling. Curr Biol, 29, 2501–2508 e3.

Butler, A., Hoffman, P., Smibert, P., Papalexi, E. & Satija, R. 2018. Integrating single-cell transcriptomic data across different conditions, technologies, and species. Nat Biotechnol, 36, 411–420.

Cano-Delgado, A., Yin, Y., Yu, C., Vafeados, D., Mora-Garcia, S., Cheng, J. C., Nam, K. H., Li, J. & Chory, J. 2004. BRL1 and BRL3 are novel brassinosteroid receptors that function in vascular differentiation in Arabidopsis. Development, 131, 5341–51.

Cattaneo, P., Graeff, M., Marhava, P. & Hardtke, C. S. 2019. Conditional effects of the epigenetic regulator JUMONJI 14 in Arabidopsis root growth. Development, 146.

Choe, S., Dilkes, B. P., Fujioka, S., Takatsuto, S., Sakurai, A. & Feldmann, K. A. 1998. The DWF4 gene of Arabidopsis encodes a cytochrome P450 that mediates multiple 22alpha-hydroxylation steps in brassinosteroid biosynthesis. Plant Cell, 10, 231–43.

Choe, S., Dilkes, B. P., Gregory, B. D., Ross, A. S., Yuan, H., Noguchi, T., Fujioka, S., Takatsuto, S., Tanaka, A., Yoshida, S., Tax, F. E. & Feldmann, K. A. 1999. The Arabidopsis dwarf1 mutant is defective in the conversion of 24-methylenecholesterol to campesterol in brassinosteroid biosynthesis. Plant Physiol, 119, 897–907.

Clark, N. M., Nolan, T. M., Wang, P., Song, G., Montes, C., Valentine, C. T., Guo, H., Sozzani, R., Yin, Y. & Walley, J. W. 2021. Integrated omics networks reveal the temporal signaling events of brassinosteroid response in Arabidopsis. Nat Commun, 12, 5858.

Crowell, H. L., Soneson, C., Germain, P. L., Calini, D., Collin, L., Raposo, C., Malhotra, D. & Robinson, M. D. 2020. Muscat detects subpopulation-specific state transitions from multi-sample multi-condition single-cell transcriptomics data. Nat Commun, 11, 6077.

Denyer, T., Ma, X., Klesen, S., Scacchi, E., Nieselt, K. & Timmermans, M. C. P. 2019. Spatiotemporal Developmental Trajectories in the Arabidopsis Root Revealed Using High-Throughput Single-Cell RNA Sequencing. Dev Cell, 48, 840–852 e5.

Dickinson, L., Yuan, W., Leblanc, C., Thomson, G., Wang, S. & Jacob, Y. 2023. Regulation of gene editing using T-DNA concatenation. Nat Plants, 9, 1398–1408.

Fabregas, N., Lozano-Elena, F., Blasco-Escamez, D., Tohge, T., Martinez-Andujar, C., Albacete, A., Osorio, S., Bustamante, M., Riechmann, J. L., Nomura, T., Yokota, T., Conesa, A., Alfocea, F. P., Fernie, A. R. & Cano-Delgado, A. I. 2018. Overexpression of the vascular brassinosteroid receptor BRL3 confers drought resistance without penalizing plant growth. Nat Commun, 9, 4680.

Fridman, Y., Elkouby, L., Holland, N., Vragovic, K., Elbaum, R. & Savaldi-Goldstein, S. 2014. Root growth is modulated by differential hormonal sensitivity in neighboring cells. Genes & Development, 28, 912–920.

Fridman, Y., Strauss, S., Horev, G., Ackerman-Lavert, M., Reiner-Benaim, A., Lane, B., Smith, R. S. & Savaldi-Goldstein, S. 2021. The root meristem is shaped by brassinosteroid control of cell geometry. Nat Plants, 7, 1475–1484.

Fukuda, H. & Ohashi-Ito, K. 2019. Vascular Essue development in plants. Curr Top Dev Biol, 131, 141–160.

Gallego-Bartolome, J., Arana, M. V., Vandenbussche, F., Zadnikova, P., Minguet, E. G., Guardiola, V., Van Der Straeten, D., Benkova, E., Alabadi, D. & Blazquez, M. A. 2011. Hierarchy of hormone action controlling apical hook development in Arabidopsis. Plant J, 67, 622–34.

Gonzalez-Garcia, M. P., Vilarrasa-Blasi, J., Zhiponova, M., Divol, F., Mora-Garcia, S., Russinova, E. & Cano-Delgado, A. I. 2011. Brassinosteroids control meristem size by promoting cell cycle progression in Arabidopsis roots. Development, 138, 849–59.

Graeff, M., Rana, S., Marhava, P., Moret, B. & Hardtke, C. S. 2020. Local and Systemic Effects of Brassinosteroid Perception in Developing Phloem. Curr Biol, 30, 1626–1638 e3.

Graeff, M., Rana, S., Wendrich, J. R., Dorier, J., Eekhout, T., Aliaga Fandino, A. C., Guex, N., Bassel, G. W., De Rybel, B. & Hardtke, C. S. 2021. A single-cell morpho-transcriptomic map of brassinosteroid action in the Arabidopsis root. Mol Plant, 14, 1985–1999.

Hacham, Y., Holland, N., Butterfield, C., Ubeda-Tomas, S., Bennett, M. J., Chory, J. & Savaldi-Goldstein, S. 2011. Brassinosteroid perception in the epidermis controls root meristem size. Development, 138, 839–48.

Hafemeister, C. & Satija, R. 2019. Normalization and variance stabilization of single-cell RNA-seq data using regularized negative binomial regression. Genome Biol, 20, 296.

Hardtke, C. S. 2023. Phloem development. New Phytol, 239, 852–867.

Hategan, L., Godza, B., Kozma-Bognar, L., Bishop, G. J. & Szekeres, M. 2014. Differential expression of the brassinosteroid receptor-encoding BRI1 gene in Arabidopsis. Planta, 239, 989–1001.

Hohmann, U., Ramakrishna, P., Wang, K., Lorenzo-Orts, L., Nicolet, J., Henschen, A., Barberon, M., Bayer, M. & Hothorn, M. 2020. Constitutive Activation of Leucine-Rich Repeat Receptor Kinase Signaling Pathways by BAK1-INTERACTING RECEPTOR-LIKE KINASE3 Chimera. Plant Cell, 32, 3311–3323.

Holzwart, E., Huerta, A. I., Glockner, N., Garnelo Gomez, B., Wanke, F., Augustin, S., Askani, J. C., Schurholz, A. K., Harter, K. & Wolf, S. 2018. BRI1 controls vascular cell fate in the Arabidopsis root through RLP44 and phytosulfokine signaling. Proc Natl Acad Sci U S A, 115, 11838–11843.

Hothorn, M., Belkhadir, Y., Dreux, M., Dabi, T., Noel, J. P., Wilson, I. A. & Chory, J. 2011. Structural basis of steroid hormone perception by the receptor kinase BRI1. Nature, 474, 467–71.

Hsu, C. W., Shahan, R., Nolan, T. M., Benfey, P. N. & Ohler, U. 2022. Protocol for fast scRNA-seq raw data processing using scKB and non-arbitrary quality control with COPILOT. STAR Protoc, 3, 101729.

Kang, Y. H., Breda, A. & Hardtke, C. S. 2017. Brassinosteroid signaling directs formative cell divisions and protophloem differentiation in Arabidopsis root meristems. Development, 144, 272–280.

Kang, Y. H. & Hardtke, C. S. 2016. Arabidopsis MAKR5 is a positive effector of BAM3-dependent CLE45 signaling. EMBO Rep, 17, 1145–54.

Kim, E. J. & Russinova, E. 2020. Brassinosteroid signalling. Curr Biol, 30, R294–R298.

Kim, H., Zhou, J., Kumar, D., Jang, G., Ryu, K. H., Sebastian, J., Miyashima, S., Helariutta, Y. & Lee, J. Y. 2020. SHORTROOT-Mediated Intercellular Signals Coordinate Phloem Development in Arabidopsis Roots. Plant Cell, 32, 1519–1535.

Koh, S. W. H., Marhava, P., Rana, S., Graf, A., Moret, B., Bassukas, A. E. L., Zourelidou, M., Kolb, M., Hammes, U. Z., Schwechheimer, C. & Hardtke, C. S. 2021. Mapping and engineering of auxin-induced plasma membrane dissociation in BRX family proteins. Plant Cell, 33, 1945–1960.

Lee, M. M. & Schiefelbein, J. 1999. WEREWOLF, a MYB-related protein in Arabidopsis, is a position-dependent regulator of epidermal cell pagerning. Cell, 99, 473–83.

Li, J. & Chory, J. 1997. A putative leucine-rich repeat receptor kinase involved in brassinosteroid signal transduction. Cell, 90, 929–38.

Li, W., Zhang, J., Tian, X., Liu, H., Ali, K., Bai, Q., Zheng, B., Wu, G. & Ren, H. 2022. Two Conserved Amino Acids Characterized in the Island Domain Are Essential for the Biological Functions of Brassinolide Receptors. Int J Mol Sci, 23.

Mccarthy, D. J., Chen, Y. & Smyth, G. K. 2012. Differential expression analysis of multifactor RNA-Seq experiments with respect to biological variation. Nucleic Acids Res, 40, 4288–97.

Mcginnis, C. S., Murrow, L. M. & Gartner, Z. J. 2019. DoubletFinder: Doublet Detection in Single-Cell RNA Sequencing Data Using Artificial Nearest Neighbors. Cell Syst, 8, 329–337 e4.

Melsted, P., Ntranos, V. & Pachter, L. 2019. The barcode, UMI, set format and BUStools. BioinformaOcs, 35, 4472–4473.

Navarro, C., Moore, J., Ott, A., Baumert, E., Mohan, A., Gill, K. S. & Sandhu, D. 2015. Evolutionary, Comparative and Functional Analyses of the Brassinosteroid Receptor Gene, BRI1, in Wheat and Its Relation to Other Plant Genomes. PLoS One, 10, e0127544.

Nolan, T. M., Vukasinovic, N., Hsu, C. W., Zhang, J., Vanhoutte, I., Shahan, R., Taylor, I. W., Greenstreet, L., Heitz, M., Afanassiev, A., Wang, P., Szekely, P., Brosnan, A., Yin, Y., Schiebinger, G., Ohler, U., Russinova, E. & Benfey, P. N. 2023. Brassinosteroid gene regulatory networks at cellular resolution in the Arabidopsis root. Science, 379, eadf4721.

Nomura, T., Kushiro, T., Yokota, T., Kamiya, Y., Bishop, G. J. & Yamaguchi, S. 2005. The last reaction producing brassinolide is catalyzed by cytochrome P-450s, CYP85A3 in tomato and CYP85A2 in Arabidopsis. J Biol Chem, 280, 17873–9.

Oh, M. H., Honey, S. H. & Tax, F. E. 2020. The Control of Cell Expansion, Cell Division, and Vascular Development by Brassinosteroids: A Historical Perspective. Int J Mol Sci, 21, 1743.

Ohashi-Ito, K., Iwamoto, K., Yamagami, A., Nakano, T. & Fukuda, H. 2023. HD-ZIP III-dependent local promotion of brassinosteroid synthesis suppresses vascular cell division in Arabidopsis root apical meristem. Proc Natl Acad Sci U S A, 120, e2216632120.

Rodriguez-Villalon, A., Gujas, B., Kang, Y. H., Breda, A. S., Cattaneo, P., Depuydt, S. & Hardtke, C. S. 2014. Molecular genetic framework for protophloem formation. Proc Natl Acad Sci U S A, 111, 11551–6.

Roh, J., Moon, J., Youn, J. H., Seo, C., Park, Y. J. & Kim, S. K. 2020. Establishment of Biosynthetic Pathways To Generate Castasterone as the Biologically Active Brassinosteroid in Brachypodium distachyon. J Agric Food Chem, 68, 3912–3923.

Savaldi-Goldstein, S., Peto, C. & Chory, J. 2007. The epidermis both drives and restricts plant shoot growth. Nature, 446, 199–202.

Sessions, A., Weigel, D. & Yanofsky, M. F. 1999. The Arabidopsis thaliana MERISTEM LAYER 1 promoter specifies epidermal expression in meristems and young primordia. Plant J, 20, 259–63.

Shahan, R., Hsu, C. W., Nolan, T. M., Cole, B. J., Taylor, I. W., Greenstreet, L., Zhang, S., Afanassiev, A., Vlot, A. H. C., Schiebinger, G., Benfey, P. N. & Ohler, U. 2022. A single-cell Arabidopsis root atlas reveals developmental trajectories in wild-type and cell identity mutants. Dev Cell, 57, 543–560 e9.

Sieburth, L. E. & Meyerowitz, E. M. 1997. Molecular dissection of the AGAMOUS control region shows that cis elements for spatial regulation are located intragenically. Plant Cell, 9, 355–65.

Stuart, T., Butler, A., Hoffman, P., Hafemeister, C., Papalexi, E., Mauck, W. M., 3rd, Hao, Y., Stoeckius, M., Smibert, P. & Satija, R. 2019. Comprehensive Integration of Single-Cell Data. Cell, 177, 1888–1902 e21.

Szekeres, M., Nemeth, K., Koncz-Kalman, Z., Mathur, J., Kauschmann, A., Altmann, T., Redei, G. P., Nagy, F., Schell, J. & Koncz, C. 1996. Brassinosteroids rescue the deficiency of CYP90, a cytochrome P450, controlling cell elongation and de-etiolation in Arabidopsis. Cell, 85, 171–82.

Tamaki, T., Oya, S., Naito, M., Ozawa, Y., Furuya, T., Saito, M., Sato, M., Wakazaki, M., Toyooka, K., Fukuda, H., Helariutta, Y. & Kondo, Y. 2020. VISUAL-CC system uncovers the role of GSK3 as an orchestrator of vascular cell type ratio in plants. Commun Biol, 3, 184.

Vragovic, K., Sela, A., Friedlander-Shani, L., Fridman, Y., Hacham, Y., Holland, N., Bartom, E., Mockler, T. C. & Savaldi-Goldstein, S. 2015. Translatome analyses capture of opposing Essue-specific brassinosteroid signals orchestrating root meristem differentiation. Proc Natl Acad Sci U S A, 112, 923–8.

Vukasinovic, N., Wang, Y., Vanhoutte, I., Fendrych, M., Guo, B., Kvasnica, M., Jiroutova, P., Oklestkova, J., Strnad, M. & Russinova, E. 2021. Local brassinosteroid biosynthesis enables optimal root growth. Nat Plants, 7, 619–632.

Wang, Z. Y., Nakano, T., Gendron, J., He, J., Chen, M., Vafeados, D., Yang, Y., Fujioka, S., Yoshida, S., Asami, T. & Chory, J. 2002. Nuclear-localized BZR1 mediates brassinosteroid-induced growth and feedback suppression of brassinosteroid biosynthesis. Dev Cell, 2, 505–13.

Yamamoto, R., Fujioka, S., Demura, T., Takatsuto, S., Yoshida, S. & Fukuda, H. 2001. Brassinosteroid levels increase drastically prior to morphogenesis of tracheary elements. Plant Physiol, 125, 556–63.

Yin, Y., Wang, Z. Y., Mora-Garcia, S., Li, J., Yoshida, S., Asami, T. & Chory, J. 2002. BES1 accumulates in the nucleus in response to brassinosteroids to regulate gene expression and promote stem elongation. Cell, 109, 181–91.

Zhang, X., Zhou, L., Qin, Y., Chen, Y., Liu, X., Wang, M., Mao, J., Zhang, J., He, Z., Liu, L. & Li, J. 2018. A Temperature-Sensitive Misfolded bri1-301 Receptor Requires Its Kinase Activity to Promote Growth. Plant Physiol, 178, 1704–1719.

Zhou, A., Wang, H., Walker, J. C. & Li, J. 2004. BRL1, a leucine-rich repeat receptor-like protein kinase, is functionally redundant with BRI1 in regulating Arabidopsis brassinosteroid signaling. Plant J, 40, 399–409.

Zhu, J. Y., Sae-Seaw, J. & Wang, Z. Y. 2013. Brassinosteroid signalling. Development, 140, 1615–20.

